# Results of a large scale study of the binding of 50 type II inhibitors to 348 kinases: The role of protein reorganization

**DOI:** 10.64898/2026.02.05.704068

**Authors:** Vardan H. Vardanyan, Allan Haldane, Howook Hwang, Dilek Coskun, Muyun Lihan, Edward B. Miller, Richard A. Friesner, Ronald M. Levy

## Abstract

Kinase family proteins constitute the second largest protein class targeted in drug development efforts, most prominently to treat cancer, but also several other diseases associated with kinase dysfunction. In this work we focus on type II kinase inhibitors which bind to the “classical” inactive conformation of the protein kinase catalytic domain where the DFG motif has a “DFG-out” orientation and the activation loop is folded. Many Tyrosine kinases (TKs) exhibit strong binding affinity with a wide spectrum of type II inhibitors while serine/threonine kinases (STKs) often bind more weakly. Recent work suggests this difference is largely due to differences in the folded to extended conformational equilibrium of the activation loop between TKs vs. STKs. The binding affinity of a type II inhibitor to its kinase target can be decomposed into a sum of two contributions: (1) the free energy cost to reorganize the protein from the active to inactive state, and (2) the binding affinity of the type II inhibitor to the inactive kinase conformation. In previous work we used a Potts statistical energy potential based on sequence co-variation to thread sequences over ensembles of active and inactive kinase structures. The threading function was used to estimate the free energy cost to reorganize kinases from the active to classical inactive conformation, and we showed that this estimator is consistent with the results of molecular dynamics free energy simulations for a small set of STKs and TKs. In the current study, we analyze the results of a large-scale study of the binding affinities of 50 type II inhibitors to 348 kinases, of which the results for 16 of the 50 type II inhibitors were reported in an earlier study (the “Davis dataset”); the binding data for the remaining 34 type II inhibitors to the panel of 348 kinases were recently obtained (the “Schrödinger dataset”). We use the Potts statistical energy model to investigate the contribution of protein reorganization to the selectivity of the large kinase panel against the set of 50 type II inhibitors, and find that protein reorganization makes a significant contribution to the selectivity. The AUC of the receiver-operator characteristic curve is ∼0.8. We report the results of an internal “blind test”, that shows how Potts threading energies can provide more accurate estimates of kinase selectivity than corresponding predictions using experimental results of small sample size. We discuss why two STK phylogenetic kinase families, STE and CMGC, appear to contain many outliers, and how to improve the ability to predict kinase selectivity with a more complete analysis of the kinase conformational landscape. We compare the performance of Potts threading for predicting binding properties of the large set of (50) Type II inhibitors to 348 kinases, with those of a sequence-based purely machine learning model, DeepDTAGen, a publicly available machine learning model that was trained on the complete Davis dataset, including both Type I and Type II kinase inhibitors. We observe that DeepDTAGen performs well on binding predictions for the 16 type II inhibitors in the Davis dataset, but performs poorly on binding predictions for the 34 type II inhibitors against 348 kinases in the Schrödinger dataset.

## I Introduction

Protein kinases are central players controlling a large number of cellular processes from cellular replication to immune response [1, 2]. Their dysregulation is implicated in many diseases; they have been studied extensively over the past thirty years [3, 4] and constitute the second largest class of druggable proteins [5, 6]. There are currently more than 120 kinase inhibitors approved or in clinical trials [7]. There have been several large-scale studies of the binding of kinase inhibitors (KIs) reported [8–11]; these papers have documented the cross-reactivity of KIs with the human kinome. The two most widely studied classes of kinase inhibitors are type I and type II. Type I inhibitors target the catalytic domain of active kinases which can bind ATP; the DFG motif is “DFG-in”, activation loop extended. The DFG motif refers to a conserved catalytic motif located at the N-terminus of a ∼20 residue long ‘activation loop’ that is highly flexible and controls the activation state of the kinase and the structure of the substrate binding surface [12]. The classical DFG-out conformation, targeted by most (but not all) type II inhibitors displays a highly reorganized activation loop that is folded away from the *α*C-helix [13]; they bind to a pocket adjacent to the ATP binding site.

Previous large-scale studies have presented the experimental binding affinities for both type I and II inhibitors, usually with a larger focus on type I inhibitors [8, 9]. In this work we focus on type II kinase inhibitors. Many Tyrosine kinases (TKs) exhibit strong binding affinity with a wide spectrum of type-II inhibitors while serine/threonine kinases (STKs) often bind more weakly. It was suggested [14, 15] that this may reflect differences between the Active /Inactive conformational equilibrium, and recent evidence strongly supports this. A widely cited large scale study in 2011 (Davis et al. [8]) reported binding affinity data for 72 kinase inhibitors with 442 kinases, including 16 type II inhibitors to 348 distinct kinases, excluding mutants and pseudo-kinases. We refer to the binding affinity data of the 16 type II inhibitors with the 348 kinases as the “Davis dataset”. We used a small subset of this data as a benchmark to develop our model for the free energy cost of kinase protein reorganization from active to inactive based on threading a Potts sequence-based statistical potential over active and inactive kinase structures [14, 16, 17]. In the current work we have substantially expanded the number of type II inhibitors for which we have binding affinity data, from 16 type II inhibitors reported in [8] to 50 type II inhibitors. Binding affinity data was purchased from Eurofins for an additional 34 type II inhibitors chosen based on the availability of structures of these inhibitors bound to kinases in the PDB. We refer to this dataset as the “Schrödinger dataset.” The combined Davis plus Schrödinger type II inhibitor binding data constitutes one of the largest datasets reporting the binding affinities of type II inhibitors to a large fraction of the kinome that we are aware of. We use the Potts statistical energy model to investigate the contribution of protein reorganization to the selectivity of the large kinase panel against the set of 50 type II inhibitors, and find that protein reorganization makes a significant contribution to the kinase selectivity.

The Davis type II inhibitor dataset includes 244 kinases (70% of the total kinase test panel) which bind to 3 or fewer type II inhibitors more strongly than 10 micromolar, among the set of 16 type II inhibitors in the Davis dataset. Apparently, based on the relatively small number (16) of the type II inhibitors included in the Davis set, a large fraction (∼2/3) of the kinases in the Davis set appear to be quite selective, but this is due to the small sample size, as only 1/3 of the kinase panel has 3 or fewer “hits” when challenged with the larger set of type II inhibitors. As described in the section below, we carried out an internal blind test to see if we could use our protein reorganization free energy model based on the Potts statistical potential to identify from the kinases that appear to be very selective based on the Davis type II dataset, those which are very selective and those which are promiscuous when challenged with the complete Davis + Schrödinger dataset, containing 50 type II inhibitors.

The structural basis for the differences in the conformational equilibrium between Active and Inactive kinase conformations for STKs vs. TKs were described in our recent work [14]. We expand upon that analysis and also discuss why two STK phylogenetic kinase families, STE and CMGC, appear to include several promiscuous kinases, despite having large threading scores (conformational equilibria strongly favoring the active state).

## II Results and discussion

### 1 Definitions of the kinase conformational landscape used in this study

The kinase conformational landscape consists of several structural basins: active DFG-in, classical DFG-out, “minimally perturbed” [14, 18], and Src-like inactive [19]. Type II inhibitors primarily target the classical DFG-out conformation, but can also bind to the “minimally perturbed” state, whereas type I inhibitors preferentially bind to the active DFG-in conformation [18, 20]. In the active DFG-in conformation, the Asp residue of the DFG motif points toward the active site, where it coordinates catalytic ions, and the activation loop adopts an extended configuration [21, 22]. In the “minimally perturbed” conformation, the DFG motif is flipped, causing the Asp residue to point away from the active site (DFG-out), while the activation loop remains extended, similar to the active state Fig. 1 (a,b). In the classical DFG-out conformation, the Asp residue of the DFG motif adopts an outward orientation, and the activation loop is folded [23], resulting in a displacement of approximately 17 Å relative to the extended loop in the active conformation Fig. 1 (c).

**Figure 1.**
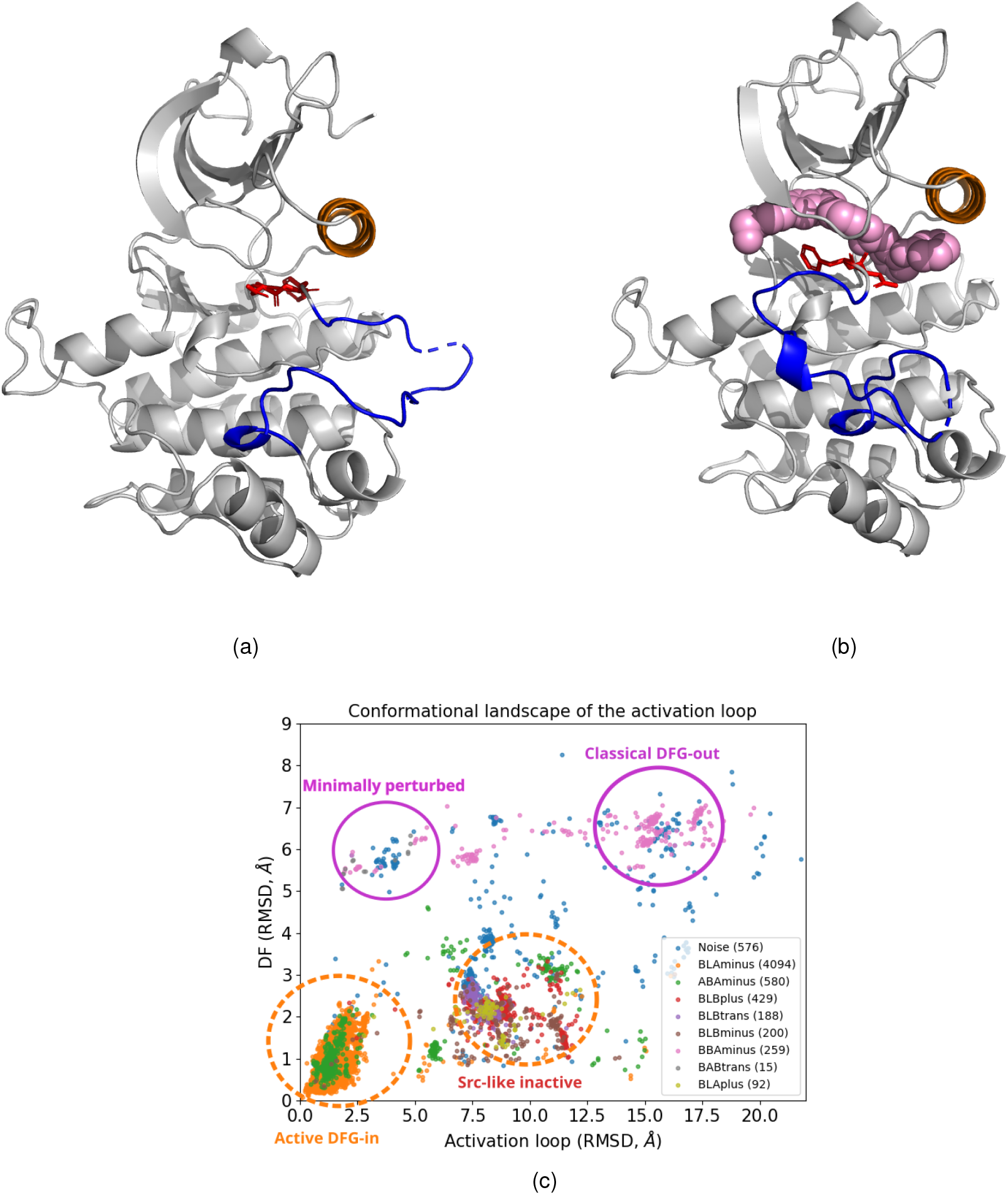
(a) INSR kinase in the active DFG-in conformation (PDB ID: 3BU3). (b) INSR kinase bound to a type II inhibitor in the classical DFG-out conformation (PDB ID: 3ETA). Color scheme: type II inhibitor in pink, DFG motif highlighted in red, activation loop shown in blue, and the *α*C-helix in brown. (c) Conformational landscape of the activation loop derived from Protein Data Bank structures classified with a scheme of [24], and plotted for two order parameters measuring the activation loop conformation as in [14].

### 2 Potts threading prediction of protein reorganization in type II inhibitor binding

The binding affinity of a type II inhibitor to its kinase target can be decomposed into a sum of two contributions: (1) the free energy cost to reorganize the protein from the active to inactive state Δ*G*_*reorg*_, and (2) the binding affinity of the type II inhibitor to the inactive kinase conformation Δ*G*_*bind*_.

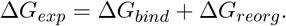

Haldane et al. [16] introduced the concept of Potts threading in which a “penalty score” (threading score Δ*T*) predicts sequence-specific conformational biases. The threading score reflects the reorganization free energy Δ*G*_*reorg*_. The Potts statistical energy model, derived from evolutionary sequence covariation [25–28], captures the conformational propensities of a given kinase sequence [29]. Threading analysis is performed between two conformational states: the active DFG-in and the classical DFG-out [14, 16, 17]. The conformational states are represented by contact maps, defined as the average contact values between residue pairs calculated from ensembles of kinase structures belonging to each respective conformation. The threading score of a sequence is then defined as the difference in Potts statistical energy between these two ensembles, providing an estimate of the energetic cost required for a kinase to reorganize from the DFG-in to the classical DFG-out state (see Eq. (2) Methods). Such large-scale conformational transitions are very difficult to observe directly [30].

Sequences with low threading scores (Δ*T*), indicating a smaller energetic cost for Δ*G*_*reorg*_, are expected to show a higher propensity for binding type II inhibitors. This relationship was previously confirmed by analyzing kinase binding assay data for 13 type II inhibitors from the Davis dataset [8,14]. Sequences with low hit rates, defined as the number of inhibitors binding to a kinase with *K*_*d*_ < 10 *µM*, were found to predominantly exhibit high threading penalties (Δ*T >* 3), indicating a reduced propensity to adopt the inactive conformation required for type II inhibitor binding.

Further analysis of kinase sequences revealed that TKs have, on average, lower threading scores than STKs, indicating a smaller energetic cost to reorganize from the active DFG-in, activation loop extended state to the classical DFG-out, activation loop folded inactive conformation. Gizzo et al. [14] constructed separate structural ensembles for both the active and inactive states of human STKs and TKs. Their analysis revealed distinctly different distributions of threading scores between the two kinase groups. To uncover the molecular basis of this evolutionary divergence in conformational landscapes, the authors employed Potts statistical energy analysis, which identified key sequence position pairs that contribute to the difference. In particular, interactions coupling the N- and C-terminal anchor residues of the activation loop to residues in the catalytic loop were found to make a substantial contribution to the observed divergence between threading score of TKs and STKs.

### 3 An expanded type II inhibitor experimental panel illustrates finite-sampling problems identifying selective kinases

In this study, we analyze a new large scale type II inhibitor experimental dataset. The Davis dataset comprises binding measurements for 16 type II inhibitors tested against 348 distinct protein kinases, obtained after excluding lipid kinases, pseudokinases, and duplicate sequence variants from the original panel [8]. In the following sections, we refer to this dataset as the Davis (16 inhibitors) dataset. The Schrödinger dataset uses the same kinase panel as the Davis dataset [8] and comprises binding affinity measurements for this kinase set against 34 type II inhibitors. These inhibitors were selected from previously published studies (Table S1) to expand the chemical space and provide broader coverage of type II binding profiles. As a result, the combined dataset includes binding affinities for 50 type II inhibitors across 348 distinct kinases.

Table 1 compares the number and percentage of kinases with hit rates ≤ 2, 3, and 4 between the Davis dataset (16 inhibitors) and the combined Davis + Schrödinger dataset (50 inhibitors). In the Davis dataset (16 inhibitors), 55% of kinases have a hit rate ≤ 2, compared with 30% in the combined Davis + Schrödinger dataset (50 inhibitors), indicating that smaller inhibitor panels tend to overestimate the number of kinases with low hit rates. When the threshold is increased to 3 and 4, the corresponding fractions rise to 66% (hit rate ≤ 3) and 73% (hit rate ≤ 4) for the 16-inhibitor panel and to 34% (hit rate ≤ 3) and 38% (hit rate ≤ 4) for the 50-inhibitor panel. The larger 50-inhibitor panel shows a more gradual increase in the percentage of kinases below each threshold, compared with the sharp rise observed for the smaller 16-inhibitor panel. These results demonstrate that expanding the number of inhibitors reduces the statistical fluctuations (including overrepresentation of kinases with an apparently high degree of selectivity) as compared to a smaller panel.

**Table 1:** Number and percentage of kinases below fixed hit rate (2,3,4) thresholds between two datasets: Davis (16 inhibitors) and Davis+Schrödinger (50 inhibitors).

| Dataset | Number of kinases<br>with hit rate $\leq 2$ | Percentage of kinases<br>with hit rate $\leq 2$ | Total number of<br>kinases |
| --- | --- | --- | --- |
| Davis (16 Inhibitors) | 190 | 55% | 348 |
| Davis+Schrödinger (50 inhibitors) | 106 | 30% | 348 |
| | Number of kinases<br>with hit rate $\leq 3$ | Percentage of kinases<br>with hit rate $\leq 3$ | |
| Davis (16 Inhibitors) | 230 | 66% | 348 |
| Davis+Schrödinger (50 inhibitors) | 120 | 34% | 348 |
| | Number of kinases<br>with hit rate $\leq 4$ | Percentage of kinases<br>with hit rate $\leq 4$ | |
| Davis (16 Inhibitors) | 253 | 73% | 348 |
| Davis+Schrödinger (50 inhibitors) | 133 | 38% | 348 |

Next, we asked whether the 16 Davis type II inhibitors represent a unique subset compared to the larger Schrödinger panel, or whether their behavior could be reproduced by randomly selecting 16 inhibitors from the expanded set. We repeatedly (10000 iterations) sampled random sets of 16 inhibitors from the combined Davis + Schrödinger datasets and compared their average hit rates with those of the original Davis panel. Table 2 summarizes the results of the randomization test. The second column reports hit rates from random selections of 16 inhibitors drawn from the entire Davis + Schrödinger panel, while the third column reports results from selections restricted to the Schrödinger set alone. In both cases, no significant differences in average hit rates were observed compared to the original Davis panel.

**Table 2:** Kinases below fixed hit rate thresholds (≤2, ≤3, ≤4), for random set of 16 inhibitors compared to Davis dataset.

|  | 50 inhibitors in combined<br>Davis + Schrödinger datasets |  | 34 inhibitors of<br>Schrödinger datasets |
| --- | --- | --- | --- |
|  | Number/percentage<br>of kinases<br>Davis 16 inhibitors | Number/percentage<br>of kinases<br>Random 16 inhibitors | Number/percentage<br>of kinases<br>Random 16 inhibitors |
| Hit rates $\leq 2$ | 190/55% | $\langle 188 \rangle / \langle 54 \rangle \%$ | $\langle 184 \rangle / \langle 53 \rangle \%$ |
| Hit rates $\leq 3$ | 244/70% | $\langle 222 \rangle / \langle 64 \rangle \%$ | $\langle 221 \rangle / \langle 63 \rangle \%$ |
| Hit rates $\leq 4$ | 264/76% | $\langle 249 \rangle / \langle 72 \rangle \%$ | $\langle 249 \rangle / \langle 72 \rangle \%$ |

### 4 Potts threading energies can provide more accurate estimates of kinase selectivity, internal blind test

To explore whether hit rates (*K*_*d*_ < 10 *µM*) derived from a small inhibitor panel can predict binding behavior in a larger screen, we selected 20 kinases Table 3 with low hit rates from the Davis dataset. These kinases were further divided into two groups based on their threading scores: 12 kinases with high threading scores and 8 kinases with low threading scores. According to our model, kinases with high threading scores are expected to incur a greater energetic cost to reorganize from the active to the inactive conformation, whereas those with low threading scores should require less reorganization free energy. Thus, although all 20 kinases exhibit low hit rates in the Davis dataset, they represent two distinct structural groups that differ in their predicted conformational flexibility.

**Table 3:**
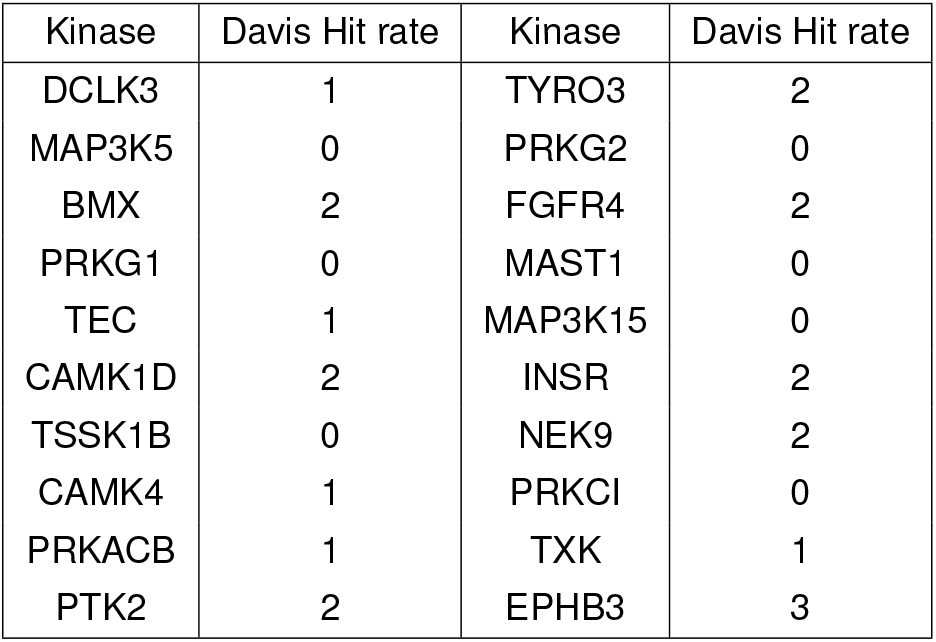
List of 20 kinases with low hit rates (*K*_*d*_ < 10 *µM*) in the Davis dataset. The table reports kinase names and corresponding hit rates from the Davis assay.

Among the 20 kinases analyzed, the Potts model predicted 12 to be selective (Table 4) and 8 to be promiscuous (Table 5), based on Potts threading scores combined with additional sequence-derived signals. These predictions were made before we received the Schrödinger dataset experimental results. Kinases with threading scores Δ*T >* 6, together with supporting sequence features, were predicted to be selective; their hit rates and strongest *K*_*d*_ values in both the Davis and Schrödinger datasets confirm this classification. One of the key sequence signals in addition to threading score was the presence of conserved paired residues R and K at positions 123 and 147, which define the two ends of the N-terminal anchor. Conversely, kinases with threading scores Δ*T <* 2 (Table 5), when combined with gatekeeper residue information [14], were consistently confirmed as promiscuous in the Schrödinger dataset.

**Table 4:** Validation of selective kinases across Davis and Schrödinger datasets.

| | Kinase | Threading<br>score $\Delta T > 6$ | Davis Hit rate<br>(16 Inh.) | Schrödinger Hit rate<br>(34 Inh.) | Davis strongest<br>$K_d$ nM | Schrödinger<br>Strongest $K_d$ nM |
| --- | --- | --- | --- | --- | --- | --- |
| 1 | PRKCI | 7.46 | 0 | 0 | 10000 | 10000 |
| 2 | PRKG2 | 7.19 | 0 | 1 | 10000 | 940 |
| 3 | CAMK4 | 7.13 | 1 | 0 | 3700 | 10000 |
| 4 | MAST1 | 6.99 | 0 | 0 | 10000 | 10000 |
| 5 | PRKG1 | 6.94 | 0 | 0 | 10000 | 10000 |
| 6 | NEK9 | 6.91 | 2 | 5 | 470 | 46 |
| 7 | DCLK3 | 6.82 | 1 | 2 | 2500 | 520 |
| 8 | CAMK1D | 6.72 | 2 | 4 | 4400 | 360 |
| 9 | MAP3K5 | 6.70 | 0 | 1 | 10000 | 2500 |
| 10 | TSSK1B | 6.64 | 0 | 0 | 10000 | 10000 |
| 11 | MAP3K15 | 6.37 | 0 | 0 | 10000 | 10000 |
| 12 | PRKACB | 6.11 | 1 | 5 | 260 | 110 |

**Table 5:** Validation of promiscuous kinases across Davis and Schrödinger datasets. L and S in paren-theses denote large and small gatekeeper residues, respectively.

| | Kinase | Threading<br>score $\Delta T < 2$ | Davis Hit rate<br>(16 Inh.) | Schrödinger Hit rate<br>(34 Inh.) | Davis strongest<br>$K_d$ nM | Schrödinger<br>Strongest $K_d$ nM |
| --- | --- | --- | --- | --- | --- | --- |
| 13 | BMX | 1.19 | 2 | 9 (S) | 200 | 6.1 |
| 14 | EPHB3 | 0.95 | 3 | 16 (S) | 85 | 71 |
| 15 | TXK | 0.75 | 1 | 11 (S) | 530 | 5.3 |
| 16 | PTK2 | 0.64 | 2 | 7 (L) | 180 | 1500 |
| 17 | TEC | 0.61 | 1 | 6 (S) | 310 | 24 |
| 18 | TYR03 | 0.19 | 2 | 7 (L) | 2.0 | 18 |
| 19 | FGFR4 | -0.33 | 2 | 8 (S) | 1400 | 11 |
| 20 | INSR | -0.58 | 2 | 5 (L) | 210 | 1300 |

The results presented in Table 4 and Table 5 motivate a broader analysis across the entire kinase panel. To this end, we plotted all 348 kinases, showing their threading scores against their hit rates to examine the overall relationship between the energetic cost of reorganization and inhibitor binding Fig. 2. The selectivity threshold for each kinase was defined with respect to the hit rate across the combined Davis + Schrödinger dataset. Kinases with a hit rate ≤ 10 were classified as selective, whereas those with a hit rate *>* 10 were classified as promiscuous. Below we test that the measurements of model performance are not strongly dependent on the threshold chosen to distinguish selective from promiscuous, as discussed below. Fig. 2 (a) shows all 348 kinases, with the CMGC and STE families in grey. Fig. 2 (b) displays the remaining kinase families, excluding CMGC and STE, for a total of 254 kinases.

**Figure 2.**
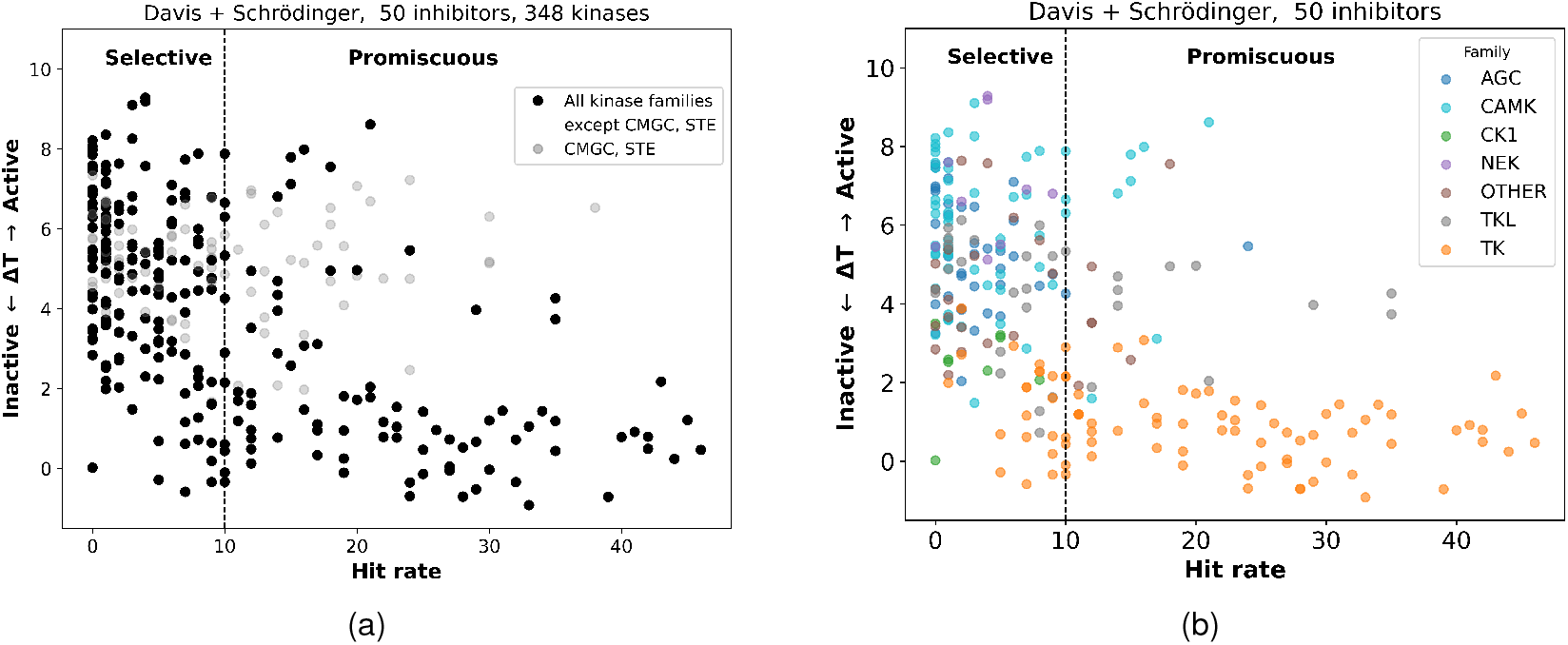
Kinase selectivity map based on threading score. (a) Scatter plot of threading score versus hit rate (*K*_*d*_ < 10 *µM*). Black points indicate the kinases included in the model selection (all families except STE and CMGC). (b) The same plot excluding STE and CMGC kinases, with remaining kinases color-coded by phylogenetic family.

Notably, the phylogenetic families belonging to STKs-including AGC, CAMK, CK1, NEK, and TKL tend to have threading scores Δ*T >* 2, whereas members of the TK family generally exhibit Δ*T <* 2. A previous study [14] reported similar differences in average threading scores between TKs and STKs, and our current analysis shows consistent behavior between threading score and hit rate, reinforcing the connection between protein reorganization to the classical inactive state targeted by type II inhibitors and selectivity.

Overall, kinases with high threading scores tend to be selective, whereas those with low threading scores are generally promiscuous, supporting the hypothesis that the threading score reflects the conformational propensity required for type II inhibitor binding. The CMGC and STE families, which deviate from this trend, are discussed in section 6 to elucidate the structural basis of their atypical behavior.

### 5 Statistical power of the Potts threading method

To further evaluate the predictive power of the threading score for identifying selective kinases, we constructed receiver operating characteristic (ROC) curves. The analysis was performed on two kinase sets: a filtered group of 254 kinases excluding the CMGC and STE families, and the complete set of 348 kinases generated using refined contact difference maps for those two kinase families, which are discussed in the following section. For these ROC curves a signal derived from the gatekeeper residue was incorporated alongside the threading score [14]. Fig. 3 (a) shows ROC curves evaluated on the kinase set excluding the CMGC and STE families for three selectivity thresholds (5, 10, and 15). The area under the curve increases from 0.80 to 0.87 as the threshold becomes more permissive. The best operating points are obtained at threading score cutoffs of approximately 3.1, 2.19, and 2.19 for the three selectivity definitions. This behavior reflects the fact that, although the positive-negative label assignment shifts with the selectivity threshold, the underlying distribution of threading scores continues to provide a consistent separation between selective and broadly active kinases. Fig. 3(b) shows the corresponding results for the complete kinase set of 348 kinases. Similar trends are observed: changing the selectivity threshold from 5 to 15 has a small effect on classifier performance, with the area under the curve rising from 0.77 to 0.80. In this case, the optimal decision threshold remains unchanged under the two stricter selectivity definitions, yielding a stable cutoff of approximately 3.7 for selectivity thresholds of 5 and 10. When the selectivity threshold is relaxed to 15, the optimal cutoff decreases to approximately 2.5, reflecting a shift in the operating point associated with the broader definition of promiscuous binding. Fig. 3(c) examines the effect of adopting a more stringent definition of a hit, reducing the affinity threshold from *K*_*d*_ < 10000 nM to *K*_*d*_ < 100 nM. Because the former represents a relatively permissive criterion for binding, this analysis probes whether tightening the hit definition alters the discriminatory power of the threading score. For the same selectivity thresholds (5, 10, and 15), the ROC curves show a consistent improvement in performance, with areas under the curve of 0.81, 0.83, and 0.84, respectively.

**Figure 3.**
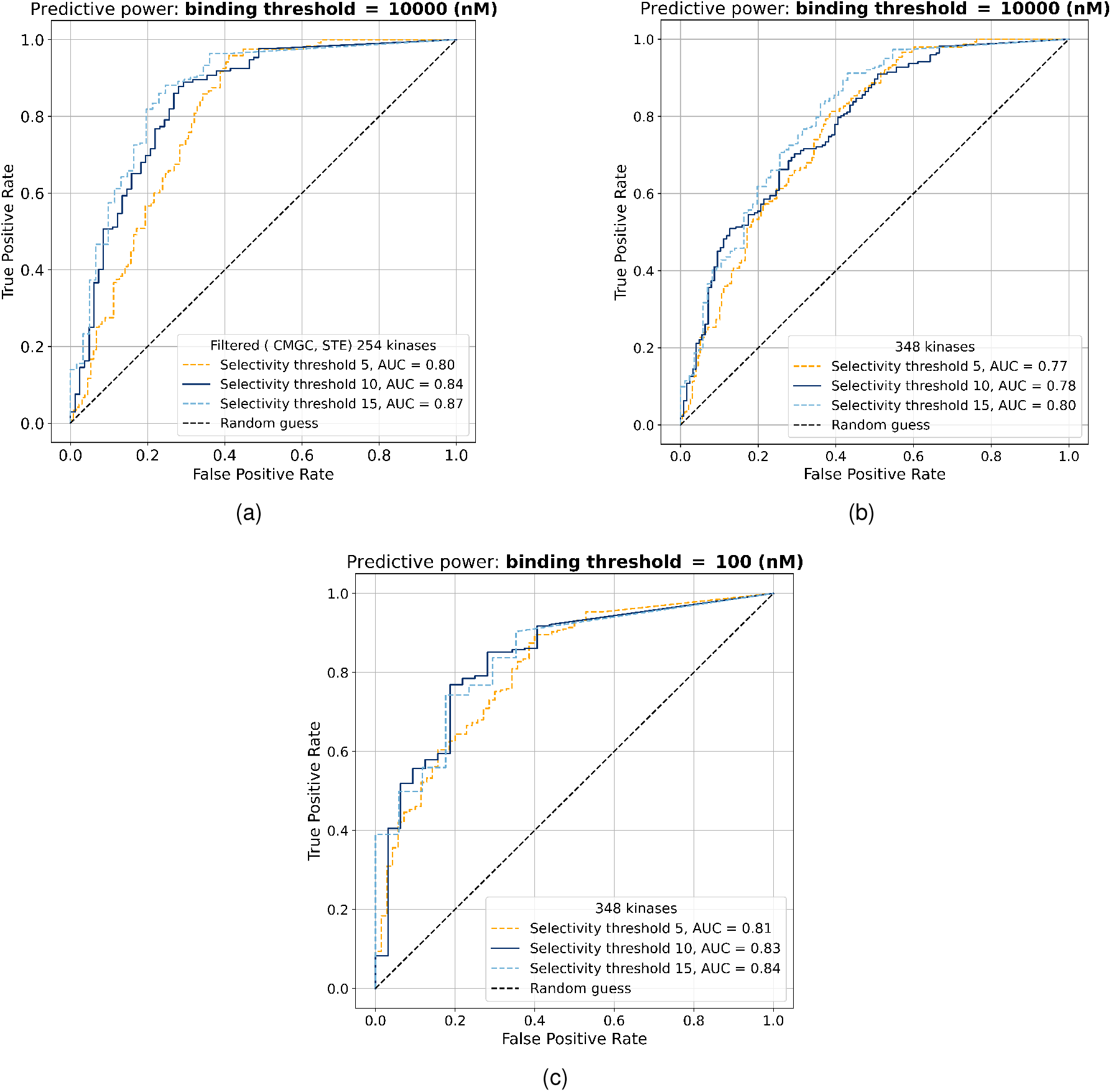
ROC curves assessing the ability of the threading score to predict kinase selectivity at selectivity thresholds of 5, 10, and 15. (a) Analysis restricted to a filtered set of 254 kinases, excluding the CMGC and STE families, with hits defined as *K*_*d*_ < 10000 nM. (b) Complete set of 348 kinases evaluated using the *K*_*d*_ < 10000 nM hit definition. (c) Complete set of 348 kinases evaluated using a more stringent hit definition of *K*_*d*_ < 100 nM.

Notably, when hits are defined at *K*_*d*_ < 100 nM, selectivity thresholds of 5, 10, or 15 may themselves become overly permissive. To address this concern, Fig S3 presents ROC curves using more stringent selectivity definitions based on hit-rate thresholds of 0, 1, 2, 3, and 4. Across this range, the area under the curve varies modestly from 0.77 to 0.81, indicating that the threading score retains comparable predictive power even under highly restrictive definitions of selectivity. Despite these small variations in ROC, the overall behavior remains consistent with the reduced kinase set: changes in the labeling scheme alter the ROC geometry and AUC values, but the threading score retains a stable and interpretable decision boundary across different selectivity regimes. These results underscore the point that the selectivity threshold itself is a somewhat arbitrary measure and varying it does not alter the underlying physical interpretation of threading score.

The threading score not only captures the previously reported trend that TKs are more promiscuous than STKs, but also reveals a finer intra-family pattern in which STKs with higher threading scores tend to be more selective than those with lower threading scores. Fig. 4(a) shows the relationship between the average threading score and average hit rate, obtained by grouping kinases into sets of 25 and computing mean values within each set to reduce noise and highlight broad trends. As the threading score decreases, the hit rate gradually increases.

**Figure 4.**
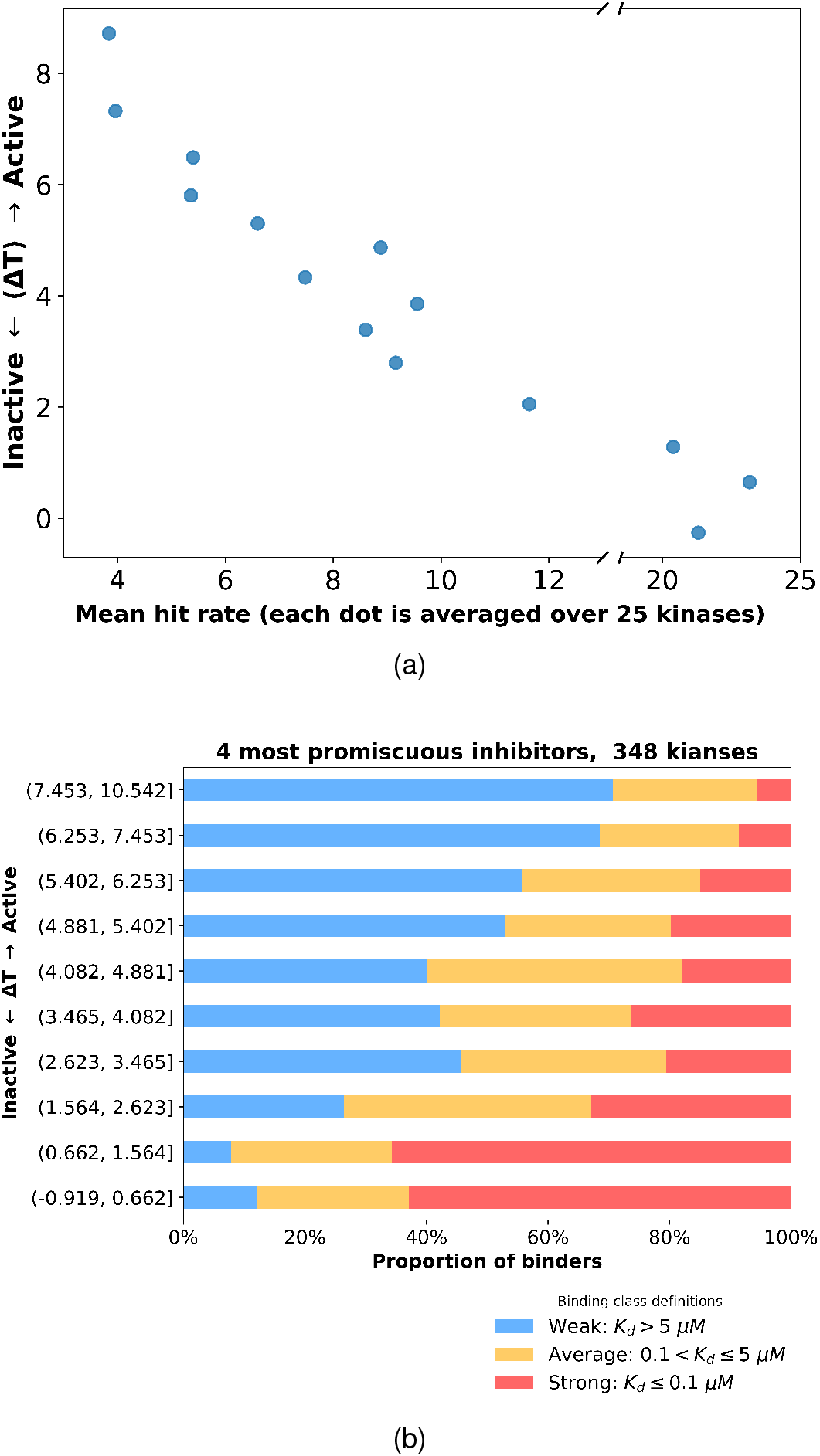
(a) Relationship between threading score and hit rate after kinases were grouped in sets of 25. Each point represents averages computed over 25 kinases, with mean hit rate on the x-axis and mean threading score on the y-axis. (b) Distribution of experimental binding affinities across threadingscore bins. Each horizontal bar represents a bin of ∼35 kinases, with colors indicating the proportion of kinases exhibiting strong (red), average (orange), and weak (blue) binding. The four most promiscuous inhibitors analyzed are Olverembatinib, Ponatinib, AST-487, and EXEL-2880|(GSK-1363089).

### 6 CMGC and STE families can bind type II inhibitor in a “minimally perturbed” conformation

In a 2022 study [14], the authors evaluated threading energies based on two contact difference maps corresponding to STKs and TKs. In the current work, we employed the same threading approach to produce the results in Fig. 2. In particular, the contact difference map for STKs is given by

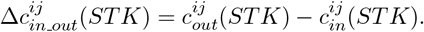

The structural ensemble of the active conformation is well characterized: the DFG motif adopts the in orientation, the activation loop is extended, and a salt bridge forms between the lysine residue of the*β*3 strand and the glutamate residue of the *α*C-helix. In contrast, the structural ensemble of inactive conformations is more heterogeneous. The activation loop for STKs can adopt either a folded position, corresponding to the classical DFG-out state (with an average shift of approximately 17 Å relative to the extended loop), a minimally perturbed position, in which the activation loop remains extended, similar to that of the active conformation, or a src-like inactive conformation which does not bind type II inhibitors.

Based on these observations we hypothesized that the reason for the different behavior of the STE and CMGC families may be due to propensity for particular inactive state. We examined which STK kinases adopt the minimally perturbed inactive conformation, characterized by a flipped DFG motif and an extended activation loop. We observed that kinases belonging to the CMGC and STE families can occupy this structural basin. In particular, this indicates that these kinases are capable of binding type II inhibitors while maintaining an extended activation loop, requiring only minimal reorganization of the protein. Thus, applying the same threading framework used for other kinase families overestimates the reorganization cost from the active to the inactive state for these two families. To address this issue, we constructed a contact difference map based on structural ensembles that include only kinases from the CMGC and STE families, which we used in all results below. Fig. S2 presents the resulting selectivity landscape, showing the threading score based organization of all 348 kinases.

### 7 Binding affinities of the kinase panel with the four most promiscuous type II inhibitors: Effects of protein reorganization

Until this point, our analysis has characterized kinase-inhibitor interactions primarily in terms of hit rates, which quantify the number of inhibitors that bind to a given kinase. This provides a coarsegrained view of the selectivity landscape. To obtain a more quantitative measure of binding strength, we next incorporated the experimentally determined dissociation constants (*K*_*d*_ values) into our analysis. Binding strength was classified based on the dissociation constant (*K*_*d*_): interactions were defined as strong for *K*_*d*_ < 100 nM, average for 100 *< K*_*d*_ ≤ 5000 nM, and weak for *K*_*d*_ *>* 5000 nM. In order to emphasize the contribution of protein reorganization (Δ*G*_*reorg*_), we focused on *K*_*d*_ patterns of the 4 most promiscuous inhibitors: Olverembatinib, Ponatinib, AST-487, and EXEL-2880| (GSK-1363089). Fig. 4 (b) shows that at high threading scores, corresponding to larger reorganization free energies (Δ*G*_*reorg*_), the fraction of kinases that bind weakly to these four promiscuous type II inhibitors, on average is much larger than the fraction of kinases that bind weakly. As the threading score decreases, the proportion of weakly binding kinases steadily declines, while the fraction of strong binders increases. This indicates that the threading score correlates with binding strength in a largely monotonic manner.

Together, the results shown in Fig. 4 establish the Potts threading score as a parameter that captures the selective-promiscuous behavior of kinases, consistent with what is observed in large inhibitor panel assays. At the coarse level of hit rates, kinases with high threading scores show low hit rates, meaning they have a low probability of binding to the inhibitors in the panel, whereas kinases with low threading scores are inhibited more frequently. When binding affinities are considered, the same trend persists: kinases with high threading scores, even when they bind, tend to do so weakly, whereas lowering the threading score is associated with a progressively higher likelihood of strong binding events.

### 8 Performance of the DeepDTAGen sequence-based Machine Learning model

In recent years, substantial effort has been devoted to predicting drug-target affinities using machine learning (ML) based approaches [31–35]. DeepDTAGen [36] is a sequence-based model and is among the most recent models reporting state-of-the-art performance across multiple evaluation benchmarks. However, the high accuracy reported by many ML based drug-target affinity predictors may, in part, reflect overfitting to training data rather than robust generalization to an unseen dataset. Given our dataset, including the Davis dataset on which DeepDTAGen was trained, and the availability of experimental binding assays for 34 previously unseen type II inhibitors against 348 kinases (the Schrödinger dataset), we viewed this as a unique opportunity to evaluate the performance of such a model.

One commonly used metric for evaluating binary classification performance, namely, whether an inhibitor binds to a kinase or not, is the precision-recall (PR) curve. DeepDTAGen reports an area under the PR curve (AUPR) of 0.77 on the complete Davis dataset, including both type I and II inhibitors, using a ground-truth binding threshold of *K*_*d*_ =100 nM. Fig. 5(a) shows the corresponding PR curve evaluated only on the subset of 16 type II inhibitors from the Davis dataset, where an increase in AUPR = 0.92 is observed. In contrast, Fig. 5(b) shows the same metric evaluated on the unseen Schrödinger dataset of 34 type II inhibitors. Here, the AUPR drops to 0.34, a substantial decrease that indicates strong overfitting.

**Figure 5.**
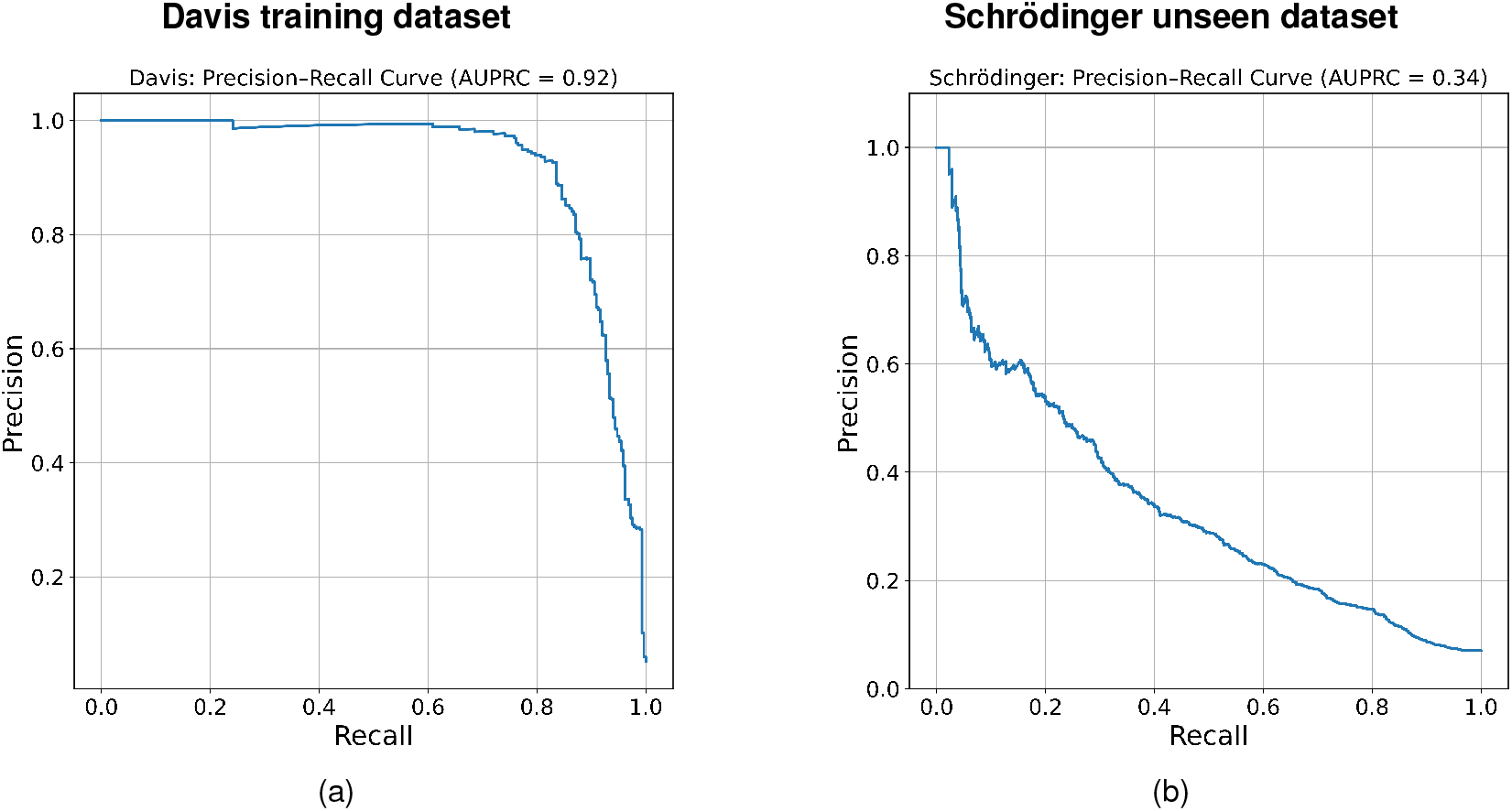
Precision-recall (PR) curve evaluation for binary binding prediction *K*_*d*_ ≤ 100 nM on (a) the Davis dataset of 16 type II inhibitors and (b) the Schrödinger dataset.

Accurate identification of strong drug-target interactions is critical for any drug-target affinity (DTA) model. To assess performance across binding strengths, we divided the binding affinities into four *K*_*d*_ ranges and evaluated DeepDTAGen within each regime Table 6. For the strongest interactions (*K*_*d*_ ≤ 10 nM), DeepDTAGen correctly identified 88 out of 120 binding events in the Davis training dataset, corresponding to a true positive rate of 73% (sensitivity or recall). In contrast, true positive rate for the same affinity range dropped to 9% when evaluated on the independent Schrödinger dataset. A similarly pronounced discrepancy between training and unseen data was observed for the *K*_*d*_ ≤ 100 nM and 100 *< K*_*d*_ ≤ 5000 nM ranges. Notably, predictive performance steadily improved for weaker binding regimes, with an acceptable true positive rate 84% achieved only for very weak binders (*K*_*d*_ *>* 5000 nM). However, approximately 81% of kinase-inhibitor pairs in the evaluated dataset fall within this weak binding regime (*K*_*d*_ *>* 5000 nM), such that the observed true positive rate (84%) represents only a modest improvement over the null expectation based on class prevalence alone.

**Table 6:** True positive rate evaluation of DeepDTAGen across four binding-affinity regimes: *K*_*d*_ ≤ 10 nM, *K*_*d*_ ≤ 100 nM, ≤ 100 *< K*_*d*_ ≤ 5000 nM, and *K*_*d*_ *>* 5000 nM. The “Experimental affinities” column reports the number of measured binding affinities in each range, the “DeepDTAGen predicted TP” column indicates the number of true positives correctly predicted by the model, and “True positive rate” is the corresponding ratio.

|  | Experimental affinities | DeepDTAGen predicted TP | True Positive Rate |
| --- | --- | --- | --- |
| **Davis (training), $K_d \leq 10$ nM** | **120** | **88** | **73%** |
| **Schrödinger, $K_d \leq 10$ nM** | **363** | **33** | **9%** |
| Davis (training), $K_d \leq 100$ nM | 275 | 221 | 80% |
| Schrödinger, $K_d \leq 100$ nM | 855 | 199 | 23% |
| Davis (training), $100 < K_d \leq 5000$ nM | 673 | 563 | 83% |
| Schrödinger, $100 < K_d \leq 5000$ nM | 1352 | 444 | 32% |
| **Davis (training), $K_d > 5000$ nM** | **4620** | **4548** | **98%** |
| **Schrödinger, $K_d > 5000$ nM** | **9625** | **8165** | **84%** |

In the previous section, we showed that threading score correlates with binding strength for the four most promiscuous inhibitors Fig. 4 (b). We focus on these inhibitors because, in the case of highly promiscuous compounds which bind to a large number of kinases (e.g. Oliverembatinib and Ponatinib each bind to more than 200 kinases in the inactive conformation with a *K*_*d*_ stronger than 10 *µM*), it is likely that the the protein-inhibitor interaction patterns with the reorganized protein are similar across many targets. Under this assumption, variations in binding affinity for highly promiscuous type II inhibitors will reflect differences in protein reorganization (Δ*G*_*reorg*_). Among the four promiscuous inhibitors considered, two originate from the Schrödinger dataset (Ponatinib and Olverembatinib), and two from the Davis dataset (AST-487 and EXEL-2880| GSK-1363089). We focus on the four inhibitors, two from the Schrödinger dataset, and two from Davis dataset to illustrate the correlation between threading score and binding strength, in a manner analogous to Fig. 4 (b). Fig. 6 (a) shows a clear relationship between threading score and experimental binding affinity for the two highly promiscuous inhibitors from the Schrödinger dataset, with higher threading scores corresponding to weaker binding and lower threading scores to stronger binding. Fig. 6 (b) maps the affinities predicted by DeepDTAGen onto kinases sorted by threading score. Notably, many experimentally observed strong binding interactions (*K*_*d*_ < 100 nM) are incorrectly predicted across both high and low threading-score regimes. The hatched regions indicate the fraction of kinases with incorrect predictions within the corresponding affinity range. Fig. 6 (b) shows that in contrast to the experimental binding affinities which are correlated with the threading scores, the binding affinities of type II inhibitors predicted by DeepDTAGen do not reflect the protein reorganization free energy (Δ*G*_*reorg*_). Fig. 6 (c) and (d) show the corresponding analyses for the Davis training dataset, in contrast to (a) and (b), which report results on the unseen Schrödinger dataset. In this case, the number of incorrect predictions is substantially reduced, with fewer strong-binding interactions misclassified across the threading-score spectrum. This difference is consistent with overfitting and further supports the conclusion that the model does not generalize well to unseen kinase-inhibitor pairs.

**Figure 6.**
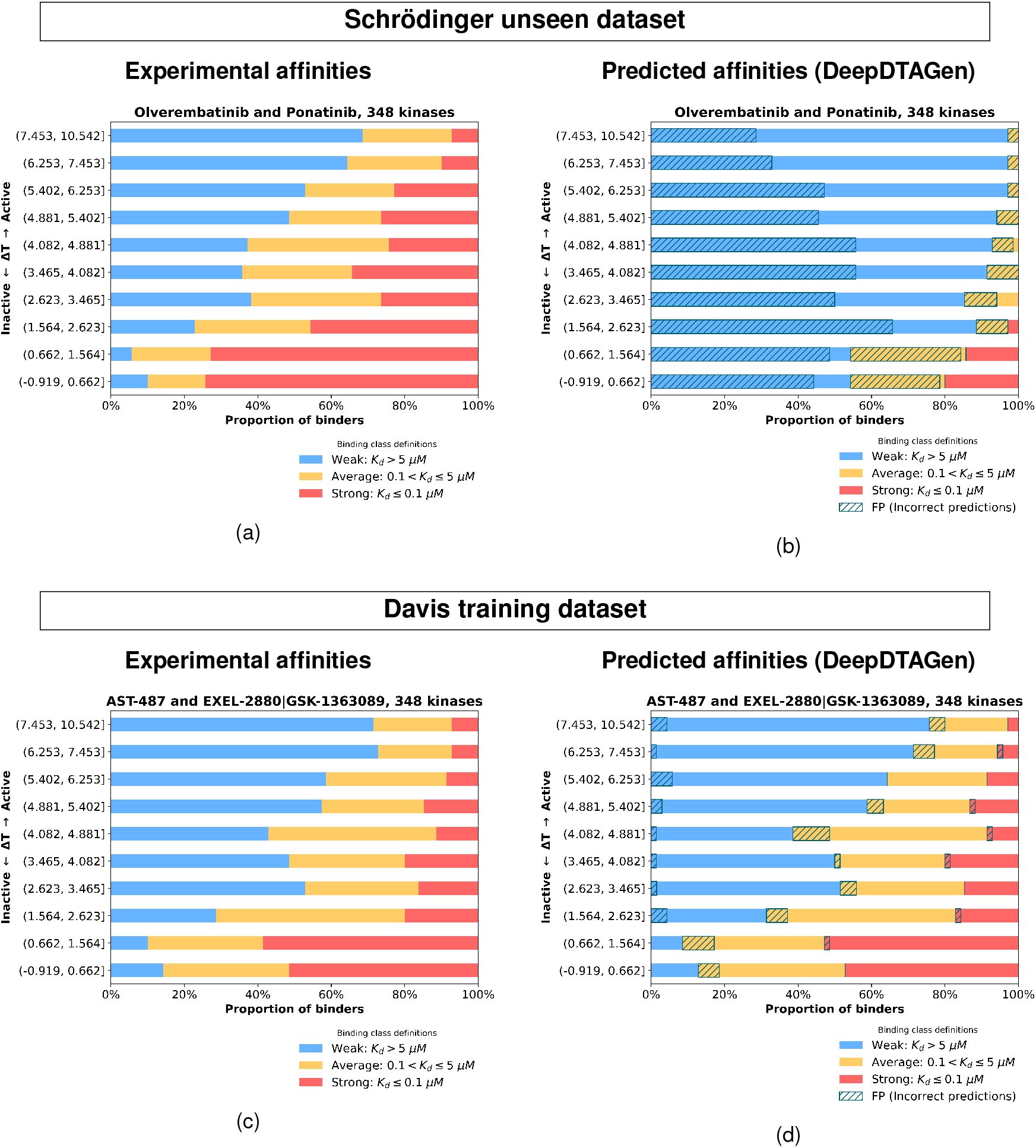
Distribution of experimental and DeepDTAGen predicted binding affinities across threading-score bins. Each horizontal bar represents a bin of ∼35 kinases, with colors indicating the proportion of kinases exhibiting strong (red), average (orange), and weak (blue) binding. (a) and (b) are the two most promiscuous inhibitors from the Schrödinger dataset (unseen dataset), showing experimental (a) and predicted (b) binding affinities, respectively. (c) and (d) show the two most promiscuous inhibitors from the Davis dataset (training dataset), with experimental (c) and predicted (d) binding affinities.

Fig. 7 shows the distribution of the 363 kinase type II inhibitor complexes with very strong binding affinities *K*_*d*_ ≤ 10 nM as a function of threading score. Consistent with the previous result, this reinforces the physical interpretation of the threading score as a proxy for the protein reorganization free energy (Δ*G*_*reorg*_), with higher threading scores corresponding to weaker binding on average and lower threading scores to stronger binding. The right panel of Fig. 7 shows the number of very strong *K*_*d*_s correctly predicted by DeepDTAGen as a function of threading score. While 29 out of the 33 correct DeepDTAGen predictions for the very strongest binders, correspond to binding to kinases with the lowest threading scores (smallest protein reorganization free energies), it is apparent that DeepDTAGen misses a large fraction (more than 90%) of strongest binders in the low-threading-score regime; in other words, it underestimates the strong binding potential of many inhibitors.

**Figure 7.**
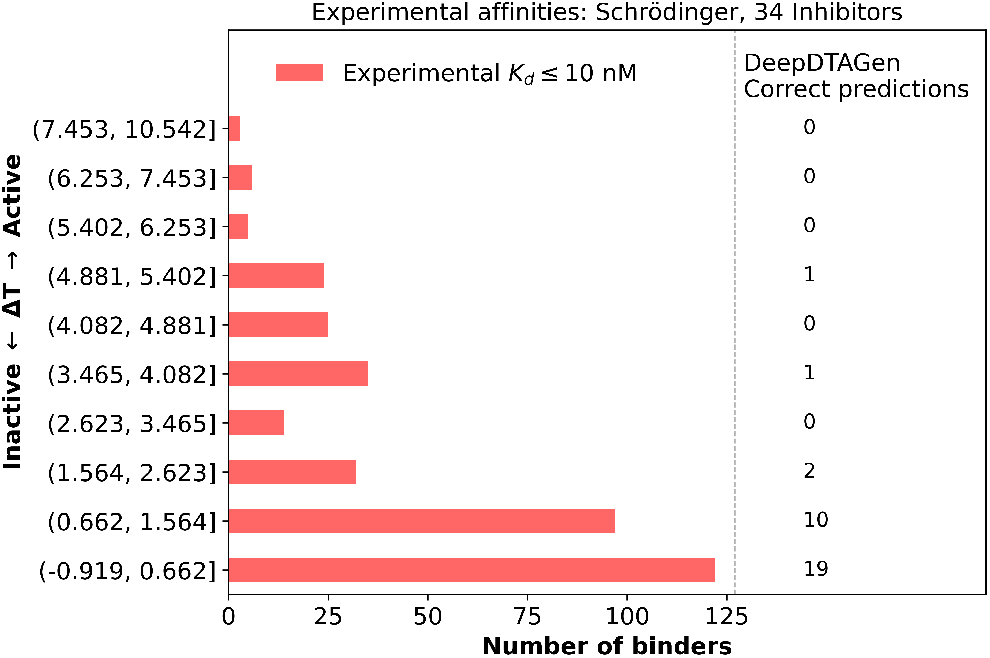
Distribution of experimental binding affinities (*K*_*d*_ ≤ 10 nM) across threading-score bins. Numbers in the right panel indicate interactions correctly predicted by DeepDTAGen as having *K*_*d*_ ≤ 10 nM.

On one hand, DeepDTAGen achieves very high scores on standard evaluation metrics when assessed on the dataset on which it was trained (the Davis dataset), but exhibits poor performance on an unseen dataset (the Schrödinger dataset), indicating substantial overfitting. On the other hand, its relatively stronger performance in the weak-binding regime, compared to the strong-binding regime, is consistent with the strong imbalance in the dataset and does not necessarily indicate enhanced discrimination of binding affinity.

DeepDTAGen is a sequence-based model, as is the Potts threading approach considered in this work, and the above comparisons therefore assess the relative performance of sequence-based methods. Another important class of machine-learning models for protein–ligand affinity prediction is structure-based. In the Supplementary Material, we evaluate a representative structure-based model, Boltz-2 [37]; its poor predictive performance in this setting provides additional context for the scope and limitations of current ML based affinity prediction approaches.

### 9 Conclusions

In this work, we analyzed the results of a large-scale study presenting the binding affinities of 50 type II inhibitors across a panel of 348 kinases. Of these inhibitors, binding data for 16 compounds were reported previously, while measurements for the remaining 34 inhibitors were obtained recently. Such large-scale profiling studies are important for quantifying kinase selectivity and for understanding how structural and phylogenetic factors govern specificity across kinase families.

We showed that many kinases appear selective when evaluated using a limited type II inhibitor panel; however, by incorporating Potts threading energies, it becomes possible to identify those kinases whose apparent selectivity arises from dataset limitations rather than intrinsic structural properties. The Potts model facilitates the prediction of kinases that are truly selective versus those likely to exhibit promiscuity when screened against a broader range of inhibitors.

We evaluated multiple receiver operating characteristic (ROC) curves by varying the selectivity threshold (hit rate ≤ 5 to hit rate ≤ 15) and found that the area under the curve (AUC) remained stable across these varying definitions. The average AUC of approximately ∼0.80 indicates that the threading score consistently provides strong predictive power for distinguishing selective from promiscuous kinases.

The availability of the Schrödinger dataset of binding affinities of 34 type II inhibitors to 348 kinases enabled a direct evaluation of the performance of one of the most recently published state-of-the-art machine-learning models for drug-target affinity (DTA) prediction, DeepDTAGen [36]. While this model performs well using standard benchmarks to evaluate its performance on the Davis type II inhibitor dataset, which formed part of the set used to train DeepDTAGen [36] its sharp performance degradation when tested on the Schrödinger type II dataset of 34 type II inhibitors binding to 348 kinases, and its systematic underestimation of strong binding interactions underscore the limitations of purely data-driven approaches and highlight the complementary value of physics-based models.

Overall, our results also show that the threading score captures more than a simple distinction between selective and promiscuous kinases. Rather than serving as a binary classifier distinguishing the selectivity of TKs from STKs, it reveals a trend across the human kinome: as the threading score increases, hit rates decrease, and a gradual shift from strong-to weak-binding inhibitors.

## III Methods

### 1 Experimental kinase inhibitor binding assays and data preparation

The Davis et al. [8] 2011 paper reports 13 type II kinase inhibitors. In that paper, the authors used the relative affinity between phosphorylated and non-phosphorylated ABL1 to determine if a compound was type II. However, this biochemical approach does not always yield definitive results. For example, the Davis et al. [8] paper classified GW-2580 as “undetermined” whereas the deposited crystal structure (PDBID: 4AT5) indicates it adopts a type 2 binding mode according to the KinCoRe kinase conformational resource [24]. This structure was released after the Davis publication. Two additional ligands, Tandutinib (MLN-518) and Linifanib (ABT-869) were reported in Davis et al. [8] as undetermined but were described as type 2 inhibitors in the literature [38, 39].

In addition to the 16 (13 reported + 1 PDB KinCoRe classified + 2 Literature declared) inhibitors identified by the Davis et al. [8] paper, we surveyed the Kincore database to collect additional type II kinase inhibitors with available structures in the PDB and which are readily purchasable from commercial vendors. This effort identified 34 type II inhibitors not reported in Davis et al. [8], resulting in a total of 50 type II kinase inhibitors utilized in this study. These compounds were then run through the Eurofin ScanMax broad kinase panel at a single concentration of 10 µM. Compounds with a reported Percent of Control (the readout of ScanMax) at 50% or smaller (smaller values indicate tighter binding) were escalated to full Kd’s with Eurofin’s KINOMEscan KdELECT Binding Assay. The complete set of ligands, their associated PDB IDs indicating type 2 binding, and the experimental assay results are provided in supplementary material.

### 2 Description of the conformational landscape

Protein kinases share a common structure consisting of an N-terminal lobe (N-lobe) and a C-terminal lobe (C-lobe) [40–42]. The N-lobe is primarily composed of β-sheets and the αC-helix, while the C-lobe is mainly α-helical [22, 43]. It contains the activation loop, a flexible segment of approximately 20–30 residues that begins with the conserved DFG (Asp-Phe-Gly) motif. Conformational changes within this loop, particularly the orientation of the DFG motif, are central to regulating kinase activity and inhibitor binding [44, 45]. The ATP-binding site is located in the cleft between the N- and C-lobes and serves as the catalytic center of the kinase. This pocket binds ATP and positions it for transfer of the γ-phosphate group to the substrate, a reaction that underlies the kinase’s enzymatic activity.

In the active conformation, the Asp residue of the DFG motif faces the ATP-binding site (DFG-in), where it helps position and stabilize two Mg^2+^ ions that are required for catalysis [46]. In this state, the activation loop adopts an open and extended configuration, allowing substrate access to the catalytic site. One of the key structural features of the active state is the Lys-Glu pair, which forms a characteristic salt bridge between the lysine residue of the β3 strand and the glutamate residue of the αC-helix. This inward positioning of the αC-helix is referred to as the αC-helix-in conformation and serves to stabilize the active state. The inactive conformation observed in many kinases corresponds to a flipped orientation of the DFG motif (DFG-out) Fig. 1 (a,b), in which the Asp residue rotates by approximately 180° relative to its position in the active state. Inactive kinases can adopt multiple inactive conformations. In these conformations, the activation loop is folded.

Gizzo et al. [14] measured the relative RMSD shifts of the DFG motif and the first five residues of the activation loop with respect to the active structure of Aurora kinase Fig. 1 (c). From these comparisons, they identified distinct structural basins corresponding to the active DFG-in, classical DFG-out, minimally perturbed, and Src-like inactive conformations. In the active DFG-in conformation, the aspartate of the DFG motif points inward toward the ATP-binding site, and the activation loop is fully extended configuration that allows substrate access. This conformation is primarily targeted by type I inhibitors. In the Src-like inactive conformation, the DFG motif remains in the same orientation as in the active DFG-in state; however, the activation loop folds, resulting in an RMSD shift of approximately 7-10 Å relative to the active conformation. In the classical DFG-out conformation, the DFG motif is flipped by 180°, resulting in an RMSD shift of about 6-8 Å relative to the active state. The activation loop is folded, resulting in an additional displacement of ∼17 Å compared to the fully extended loop observed in the active conformation Fig. 1 (b). This conformation is primarily targeted by type II inhibitors, although such inhibitors are not restricted to the classical DFG-out state. In the minimally perturbed conformation, the DFG motif is flipped by approximately 180°, with the aspartate residue pointing away from the ATP-binding site. However, unlike the classical DFG-out state, the activation loop remains fully extended. In the following sections, we present evidence that the CMGC and STE phylogenetic families are capable of binding type II inhibitors while adopting this conformation.

### 3 PDB dataset

X-ray crystal structures of eukaryotic protein kinases were downloaded from the Protein Data Bank (PDB) (http://rcsb.org) on July 30, 2020. Protein sequences from 7919 chains of 6805 PDB entries were extracted by parsing the SEQRES records. PDB structures were converted into matrices of binary contacts, where a value of 1 indicates a contact between residues and 0 denotes no contact. Residues *i* and *j* were considered to be in contact if the distance between them was less than 6 Å (*r*_*ij*_ < 6 Å). The contact frequency 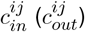 was calculated as

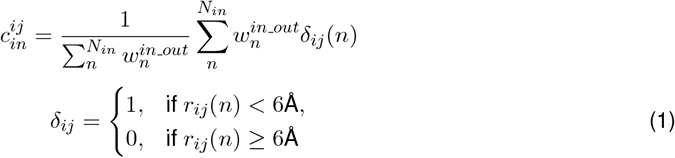

where *N*_*in*_ is the number of structures in the active cluster, and the weights were computed as 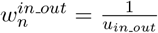 with *u*_*in*_*_*_*out*_ representing the number of occurrences of the same kinase within either cluster. This weighting scheme down-weights the contributions of kinases that are overrepresented in the PDB.

### 4 Potts model and threading

The Potts Hamiltonian was constructed from a multiple sequence alignment (MSA) of protein kinase catalytic domains, as previously described [14,47]. The probability of observing a sequence *S* is given by

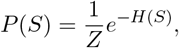

with normalization constant 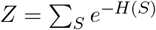.

*H*(*S*) denotes the statistical energy, defined as Potts Hamiltonian

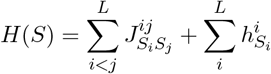

where *L* is the number of columns in the MSA (*L* = 259. In this model, the parameters 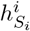 represent the statistical energy contribution of residue *S*_*i*_ at position *i*, and 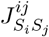 represents the pairwise coupling energy between residues *S*_*i*_ and *S*_*j*_ at positions *i* and *j*, respectively.

The threaded energy penalty, Δ*T* (*S*), quantifying the cost for sequence *S* to undergo the conformational transition from active to inactive, was calculated using contact frequency differences between the two conformational ensembles [16]:

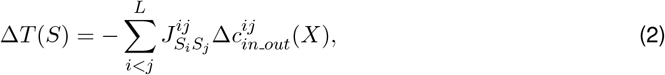

Here, *X* denotes the kinase class (TK or STK) to which sequence *S* belongs, and 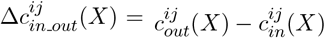 represents the difference in contact frequencies between active and inactive conformations, computed from other sequences within the same class *X*.

### 5 Dataset preparation using DeepDTAGen predicted binding affinities

The DeepDTAGen framework consists of multiple prediction modules; in this work, we used the Fully Connected Module to predict drug–target binding affinities. The pretrained DeepDTAGen model was downloaded directly from the official GitHub repository (https://github.com/CSUBioGroup/DeepDTAGen). Predicted binding affinities were generated for drug–target pairs using this pretrained model. Protein sequences used as model inputs were taken directly from the file provided with the repository, ensuring consistency with the pretrained model configuration. Small-molecule structures were represented using canonicalized isomeric SMILES (isoSMILES) generated from the original compound SMILES strings. The pretrained DeepDTAGen model was validated by comparing predicted affinities with experimentally measured binding affinities from the Davis dataset.

## Supplementary Information

The Supplementary Information presents an evaluation of the Boltz-2 structure-based machine-learning model, a complete list of all 50 type II inhibitors with their number of hits and year of development, and analysis of the selectivity landscape, including ROC curves across selectivity thresholds of 0, 1, 2, 3, 4. Binding results from 442 kinase assays for 34 type II inhibitors in the Schrödinger dataset, together with binding data for an additional 16 type II inhibitors from the Davis dataset, are provided as an Excel spreadsheet.

## 6 Acknowledgments

This research was supported by National Institutes of Health Grant No. R35 GM132090 (R.M.L) and by NIH Computer Equipment Grant No. OD020095. This work was supported in part by the Gates Foundation INV-076523. The conclusions and opinions expressed in this work are those of the author(s) alone and shall not be attributed to the Foundation. Under the grant conditions of the Foundation, a Creative Commons Attribution 4.0 License has already been assigned to the Author Accepted Manuscript version that might arise from this submission. Please note works submitted as a preprint have not undergone a peer review process.

