## Supplementary Information for "Results of a large scale study of the binding of 50 type II inhibitors to 348 kinases: The role of protein reorganization"

### Supplementary Material

#### Performance of the Boltz-2 structure-based Machine Learning model

Protein-ligand interaction prediction models can be broadly categorized into two classes: sequence- based models and structure-based models [1]. In section 8, we introduced the sequence-based framework DeepDTA-Gen [2] and evaluated its predictive power on the Davis training dataset and the Schrödinger unseen dataset. In the present section, we shift focus to structure-based approaches and examine Boltz-2 [3], a co-folding model designed to predict three-dimensional molecular structures and binding affinities of protein-ligand complexes, as well as complexes involving DNA and RNA.

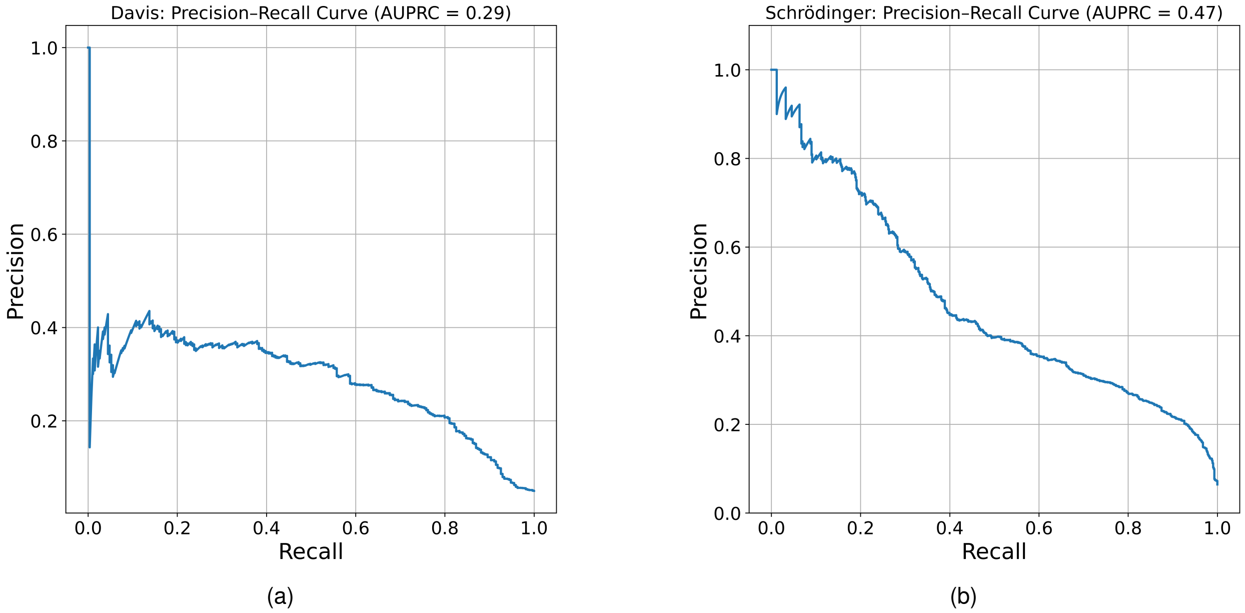


Figure 1: Precision-Recall curve evaluation for binary binding prediction $K_d < 100$ nM on a) the 16 type II inhibitors from Davis dataset and b) 34 type II inhibitors from the Schrödinger dataset.

The performance of Boltz-2 was evaluated using 16 type II inhibitors from the Davis dataset Fig. 1 (a) and 34 Type II inhibitors from the Schrödinger dataset Fig. 1(b), following the same evaluation protocol used for DeepDTAGen. Boltz-2 achieved an AUPRC of 0.28 on the Davis dataset, compared to 0.46 on the Schrödinger dataset. These results indicate poor overall predictive performance, with limited discrimination beyond baseline expectations across both datasets.

#### Inhibitors

Table S1: List of 50 type II kinase inhibitors. The table provides the inhibitor index, name, Number of hits (across 348-kinase panel), year of publication, and the corresponding reference. Imatinib, the first developed inhibitor in this list (highlighted in the 38th row), is included as a reference point.

| # | Inhibitor | Number of hits | Year | Citation |
|---|---|---|---|---|
| 1 | Olverembatinib | 229 | 2013 | [4] |
| 2 | Ponatinib | 224 | 2010 | [5,6] |
| 3 | AST-487 | 206 | 2007 | [7] |

Continued on next page

Table S1

| # | Inhibitor | Number of hits | Year | Citation |
|---|---|---|---|---|
| 4 | EXEL-2880\|GSK-1363089 | 188 | 2005 | [8] |
| 5 | Rebastinib | 181 | 2011 | [9] |
| 6 | NG25 | 155 | 2014 | [10] |
| 7 | RIPK1-IN-4 | 145 | 2013 | [11] |
| 8 | Golvatinib | 133 | 2009 | [12] |
| 9 | LY3009120 | 123 | 2015 | [13] |
| 10 | MAPK13-IN-1 | 111 | 2012 | [14] |
| 11 | Tovorafenib | 109 | 2017 | [15] |
| 12 | Merestinib | 101 | 2013 | [16] |
| 13 | Sorafenib | 100 | 2001 | [17] |
| 14 | Linifanib (ABT-869) | 88 | 2007 | [18] |
| 15 | PF-6683324 | 77 | 2018 | [19] |
| 16 | PLX-4720 | 74 | 2008 | [20] |
| 17 | Bafetinib | 71 | 2007 | [21] |
| 18 | BIRB-796 | 71 | 2002 | [22] |
| 19 | Pexidartinib | 68 | 2004 | [23] |
| 20 | DDR1-IN-1 | 66 | 2013 | [24] |
| 21 | NVP-BHG712 | 65 | 2016 | [25] |
| 22 | CHIR-265\|RAF-265 | 65 | 2006 | [26, 27] |
| 23 | TAK-632 | 65 | 2013 | [28] |
| 24 | AZD-1152HQPA | 62 | 2007 | [29, 30] |
| 25 | Ki-20227 | 62 | 2006 | [31] |
| 26 | DDR Inhibitor | 60 | 2015 | [32] |
| 27 | Nilotinib | 52 | 2007 | [33] |
| 28 | GNE-9815 | 51 | 2021 | [34] |
| 29 | Belvarafenib | 48 | 2021 | [35] |
| 30 | p38-a MAPK-IN-1 | 45 | 2009 | [36] |
| 31 | AMG-706 | 42 | 2007 | [37] |
| 32 | AWL-II-38.3 | 40 | 2009 | [38] |
| 33 | AAL993 | 39 | 2015 | [39] |
| 34 | ALW-II-49-7 | 39 | 2009 | [38] |
| 35 | MLN-518 | 37 | 2002 | [40] |
| 36 | AC220 | 36 | 2009 | [41] |
| 37 | RAF709 | 32 | 2017 | [42] |

Continued on next page

Table S1

| # | Inhibitor | Number of hits | Year | Citation |
|---|---|---|---|---|
| 38 | NVP-BHG712 isomer | 32 | 2016 | [25] |
| 39 | Imatinib | 29 | 1996 | [43, 44] |
| 40 | PDGFRa kinase inhibitor 1 | 28 | 2017 | [45] |
| 41 | GSK2334470 | 23 | 2011 | [46] |
| 42 | AB-1010 | 23 | 2008 | [47] |
| 43 | Exarafenib | 21 | 2024 | [48] |
| 44 | B-Raf IN 1 | 21 | 2009 | [49] |
| 45 | Naporafenib | 21 | 2019 | [50] |
| 46 | SR-318 | 14 | 2019 | [51] |
| 47 | DDR1-IN-4 | 9 | 2018 | [52] |
| 48 | GW-2580 | 4 | 2006 | - |
| 49 | MP7 | 3 | 2011 | [53] |
| 50 | TH470 | 3 | 2022 | [54] |

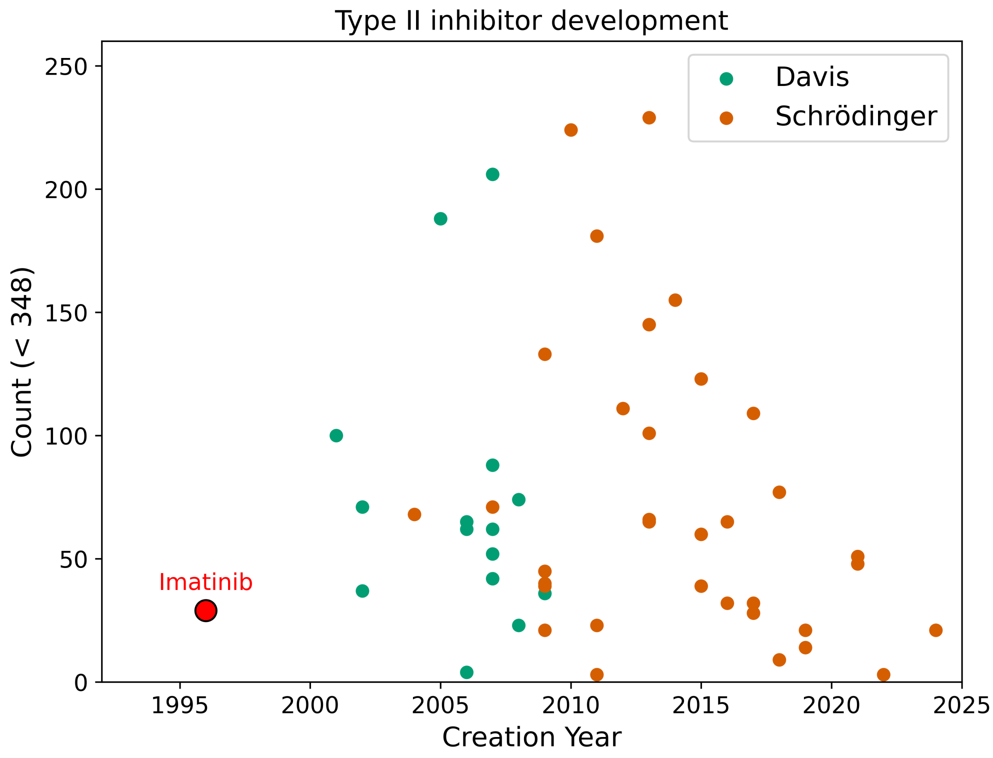


Figure S1: Selectivity of type II kinase inhibitors over time. The x-axis indicates the year of development, and the y-axis shows inhibitor selectivity (number of kinases bound out of a 348-kinase panel). Imatinib, the first type II inhibitor developed, is shown explicitly. Data points are color-coded: green for inhibitors reported in Davis et al. and brown for inhibitors reported in Schrödinger.

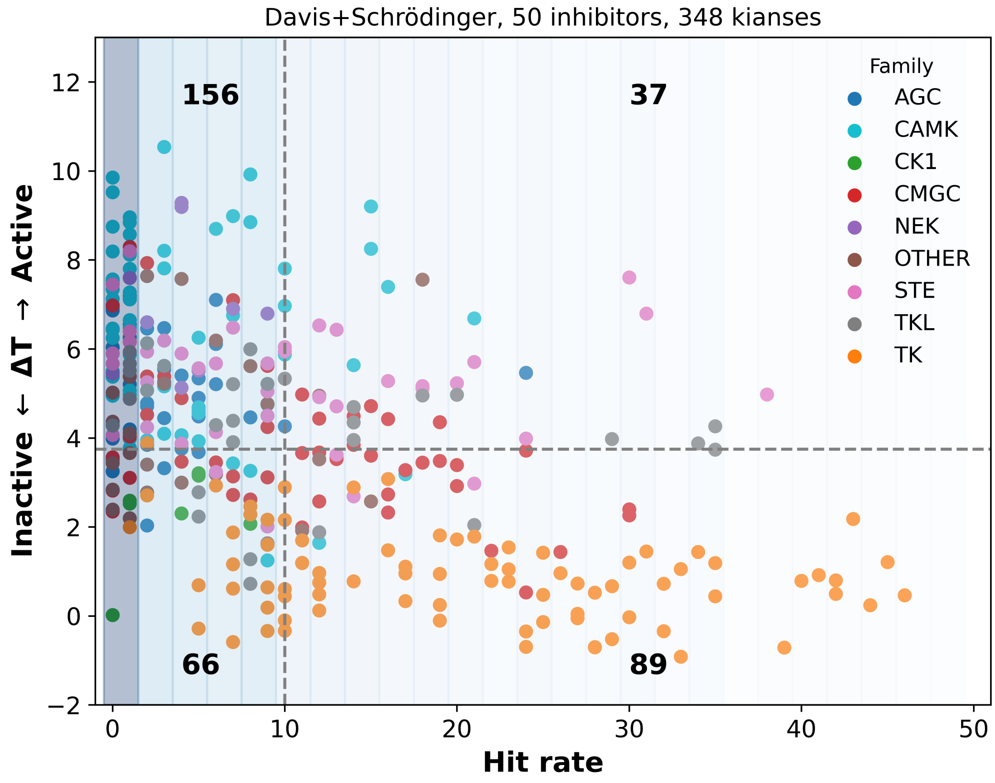


Figure S2: Selectivity landscape constructed using the new contact difference map (CDM) for three kinase families: CMGC, STE and CAMK. The vertical dashed line indicates the selectivity barrier, separating selective kinases from promiscuous ones. The horizontal dashed line represents the optimal threading-score threshold determined from the ROC analysis (Fig. 3), distinguishing kinases predicted to be promiscuous from those classified as selective. Numbers within each square denote the count of kinases falling into that region. Vertical color bars indicate the density of kinases per bin, with darker shading corresponding to bins containing more kinases.

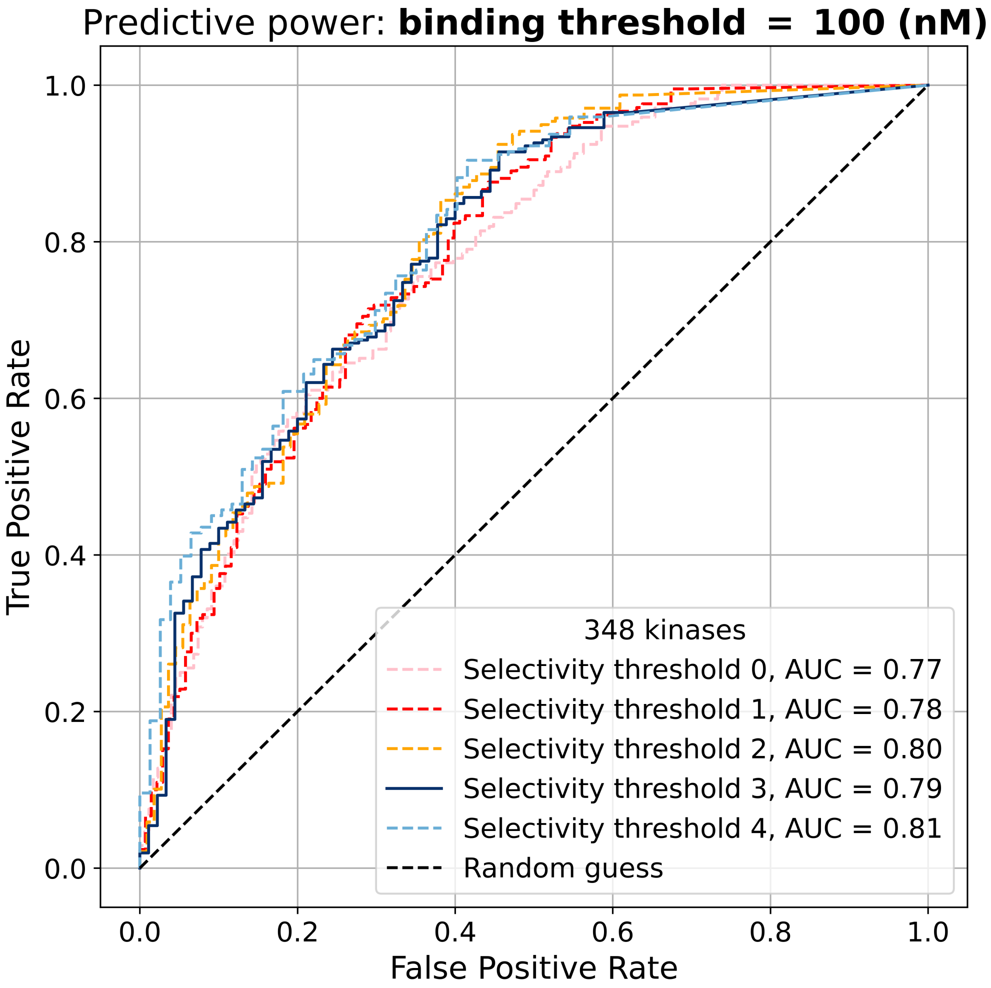


Figure S3: ROC curves assessing the ability of the threading score to predict kinase selectivity at selectivity thresholds of 0,1,2,3 and 4. Complete set of 348 kinases evaluated using the $K_d < 100$ nM hit definition.

[6] Wei Sheng Huang, Chester A. Metcalf, Raji Sundaramoorthi, Yihan Wang, Dong Zou, R. Mathew Thomas, Xiaotian Zhu, Lisi Cai, David Wen, Shuangying Liu, Jan Romero, Jiwei Qi, Ingrid Chen, Geetha Banda, Scott P. Lentini, Sasmita Das, Qihong Xu, Jeff Keats, Frank Wang, Scott Wardwell, Yaoyu Ning, Joseph T. Snodgrass, Marc I. Broudy, Karin Russian, Tianjun Zhou, Lois Commodore, Narayana I. Narasimhan, Qurish K. Mohemmad, John Iuliucci, Victor M. Rivera, David C. Dalgarno, Tomi K. Sawyer, Tim Clackson, and William C. Shakespeare. Discovery of 3-[2-(imidazo[1,2- b ]pyridazin-3-yl)ethynyl]-4-methyl- n -{4-[(4-methylpiperazin-1-yl)methyl]-3-(trifluoromethyl)phenyl}benzamide (ap24534), a potent, orally active pan-inhibitor of breakpoint cluster region-abelson (bcr-abl) kinase including the t315i gatekeeper mutant. *Journal of Medicinal Chemistry*, 53:4701–4719, 6 2010.

[7] Nagako Akeno-Stuart, Michelle Croyle, Jeffrey A. Knauf, Roberta Malaguarnera, Donata Vitagliano, Massimo Santoro, Christine Stephan, Konstantina Grosios, Markus Wartmann, Robert Cozens, Giorgio Caravatti, Doriano Fabbro, Heidi A. Lane, and James A. Fagin. The ret kinase inhibitor nvp-ast487 blocks growth and calcitonin gene expression through distinct mechanisms in medullary thyroid cancer cells. *Cancer Research*, 67:6956–6964, 7 2007.

[8] Fawn Qian, Stefan Engst, Kyoko Yamaguchi, Yu Peiwen, Kwang Ai Won, Lillian Mock, Tracy Lou, Jenny Tan, Connie Li, Danny Tam, Julie Lougheed, F. Michael Yakes, Frauke Bentzien, Xu Wei, Tal Zaks, Richard Wooster, Joel Greshock, and Alison H. Joly. Inhibition of tumor cell growth, invasion, and metastasis by exel-2880 (xl880, gsk1363089), a novel inhibitor of hgf and vegf receptor tyrosine kinases. *Cancer Research*, 69:8009–8016, 10 2009.

[9] Wayne W. Chan, Scott C. Wise, Michael D. Kaufman, Yu Mi Ahn, Carol L. Ensinger, Torsten Haack, Molly M. Hood, Jennifer Jones, John W. Lord, Wei Ping Lu, David Miller, William C. Patt, Bryan D. Smith, Peter A. Petillo, Thomas J. Rutkoski, Hanumaiah Telikepalli, Lakshminarayana Vogeti, Tony Yao, Lawrence Chun, Robin Clark, Peter Evangelista, L. Cristina Gavrilescu, Katherine Lazarides, Virginia M. Zaleskas, Lance J. Stewart, Richard A. Van Etten, and Daniel L. Flynn. Conformational control inhibition of the bcr-abl1 tyrosine kinase, including the gatekeeper t315i mutant, by the switch-control inhibitor dcc-2036. *Cancer Cell*, 19:556–568, 2011.

[10] Li Tan, Tyzoon Nomanbhoy, Deepak Gurbani, Matthew Patricelli, John Hunter, Jiefei Geng, Lina Herhaus, Jianming Zhang, Eduardo Pauls, Youngjin Ham, et al. Discovery of type ii inhibitors of tgf$\beta$-activated kinase 1 (tak1) and mitogen-activated protein kinase kinase kinase kinase 2 (map4k2). *Journal of medicinal chemistry*, 58(1):183–196, 2015.

[11] Philip A. Harris, Deepak Bandyopadhyay, Scott B. Berger, Nino Campobasso, Carol A. Capriotti, Julie A. Cox, Lauren Dare, Joshua N. Finger, Sandra J. Hoffman, Kirsten M. Kahler, Ruth Lehr, John D. Lich, Rakesh Nagilla, Robert T. Nolte, Michael T. Ouellette, Christina S. Pao, Michelle C. Schaeffer, Angela Smallwood, Helen H. Sun, Barbara A. Swift, Rachel D. Totoritis, Paris Ward, Robert W. Marquis, John Bertin, and Peter J. Gough. Discovery of small molecule rip1 kinase inhibitors for the treatment of pathologies associated with necroptosis. *ACS Medicinal Chemistry Letters*, 4:1238–1243, 12 2013.

[12] Takayuki Nakagawa, Osamu Tohyama, Atsumi Yamaguchi, Tomohiro Matsushima, Keiko Takahashi, Setsuo Funasaka, Shuji Shirotori, Makoto Asada, and Hiroshi Obaishi. E7050: A dual c-met and vegfr-2 tyrosine kinase inhibitor promotes tumor regression and prolongs survival in mouse xenograft models. *Cancer Science*, 101:210–215, 1 2010.

[13] James R. Henry, Michael D. Kaufman, Sheng-Bin Peng, Yu Mi Ahn, Timothy M. Caldwell, Lakshminarayana Vogeti, Hanumaiah Telikepalli, Wei-Ping Lu, Molly M. Hood, Thomas J. Rutkoski, Bryan D. Smith, Subha Vogeti, David Miller, Scott C. Wise, Lawrence Chun, Xiaoyi Zhang, Youyan Zhang, Lisa Kays, Philip A. Hipskind, Aaron D. Wrobleski, Karen L. Lobb, Julia M. Clay, Jeffrey D. Cohen, Jennie L. Walgren, Denis McCann, Phenil Patel, David K. Clawson, Sherry Guo, Danalyn Manglicmot, Chris Groshong, Cheyenne Logan, James J. Starling, and Daniel L. Flynn. Discovery of 1-(3,3-dimethylbutyl)-3-(2-fluoro-4-methyl-5-(7-methyl-2-(methylamino)pyrido[2,3-d]pyrimidin-6-yl)phenyl)urea (ly3009120) as a pan-raf inhibitor with minimal paradoxical activation and activity against braf or ras mutant tumor cells. *Journal of Medicinal Chemistry*, 58(10):4165–4179, 2015.

[14] Yael G. Alevy, Anand C. Patel, Arthur G. Romero, Dhara A. Patel, Jennifer Tucker, William T. Roswit, Chantel A. Miller, Richard F. Heier, Derek E. Byers, Tom J. Brett, and Michael J. Holtzman. Il-13-induced airway mucus production is attenuated by mapk13 inhibition. *Journal of Clinical Investigation*, 122:4555–4568, 12 2012.

[15] Yu Sun, John A. Alberta, Catherine Pilarz, David Calligaris, Emily J. Chadwick, Shakti H. Ramkissoon, Lori A. Ramkissoon, Veronica Matia Garcia, Emanuele Mazzola, Liliana Goumnerova, Michael Kane, Zhan Yao, Mark W. Kieran, Keith L. Ligon, William C. Hahn, Levi A. Garraway, Neal Rosen, Nathanael S. Gray, Nathalie Y. Agar, Sara J. Buhrlage, Rosalind A. Segal, and Charles D. Stiles. A brain-penetrant raf dimer antagonist for the noncanonical braf oncoprotein of pediatric low-grade astrocytomas. *Neuro-Oncology*, 19:774–785, 2017.

[16] S. Betty Yan, Victoria L. Peek, Rose Ajamie, Sean G. Buchanan, Jeremy R. Graff, Steven A. Heidler, Yu Hua Hui, Karen L. Huss, Bruce W. Konicek, Jason R. Manro, Chuan Shih, Julie A. Stewart, Trent R. Stewart, Stephanie L. Stout, Mark T. Uhlik, Suzane L. Um, Yong Wang, Wenjuan Wu, Lei Yan, Wei J. Yang, Boyu Zhong, and Richard A. Walgren. Ly2801653 is an orally bioavailable multi-kinase inhibitor with potent activity against met, mst1r, and other oncoproteins, and displays anti-tumor activities in mouse xenograft models. *Investigational New Drugs*, 31:833–844, 8 2013.

[17] J F Lyons, S Wilhelm, B Hibner, and G Bollag. Discovery of a novel raf kinase inhibitor ras epidemiology. Technical report, 2001.

[18] Yujia Dai, Kresna Hartandi, Zhiqin Ji, Asma A Ahmed, Daniel H Albert, Joy L Bauch, Jennifer J Bouska, Peter F Bousquet, George A Cunha, Keith B Glaser, et al. Discovery of n-(4-(3-amino-1 h-indazol-4-yl) phenyl)-n '-(2-fluoro-5-methylphenyl) urea (abt-869), a 3-aminoindazole-based orally active multitargeted receptor tyrosine kinase inhibitor. *Journal of medicinal chemistry*, 50(7):1584–1597, 2007.

[19] Luping Qiu, Kymberly Levine, Ketan S. Gajiwala, Ciarán N. Cronin, Asako Nagata, Eric Johnson, Michelle Kraus, John Tatlock, Robert Kania, Timothy Foley, and Shaoxian Sun. Small molecule inhibitors reveal ptk6 kinase is not an oncogenic driver in breast cancers. *PLoS ONE*, 13, 6 2018.

[26] Clara Montagut and Jeff Settleman. Targeting the raf-mek-erk pathway in cancer therapy, 10 2009.

[27] Teresa E. Williams, Sharadha Subramanian, Joelle Verhagen, Christopher M. McBride, Abran Costales, Leonard Sung, William Antonios-Mccrea, Maureen McKenna, Alicia K. Louie, Savithri Ramurthy, Barry Levine, Cynthia M. Shafer, Timothy MacHajewski, Paul A. Renhowe, Brent A. Appleton, Payman Amiri, James Chou, Darrin Stuart, Kimberly Aardalen, and Daniel Poon. Discovery of raf265: A potent mut-b-raf inhibitor for the treatment of metastatic melanoma. *ACS Medicinal Chemistry Letters*, 6:961–965, 9 2015.

[28] Masanori Okaniwa, Masaaki Hirose, Takeo Arita, Masato Yabuki, Akito Nakamura, Terufumi Takagi, Tomohiro Kawamoto, Noriko Uchiyama, Akihiko Sumita, Shunichirou Tsutsumi, Tsuneaki Tottori, Yoshitaka Inui, Bi Ching Sang, Jason Yano, Kathleen Aertgeerts, Sei Yoshida, and Tomoyasu Ishikawa. Discovery of a selective kinase inhibitor (tak-632) targeting pan-raf inhibition: Design, synthesis, and biological evaluation of c -7-substituted 1,3-benzothiazole derivatives. *Journal of Medicinal Chemistry*, 56:6478–6494, 8 2013.

[29] Robert W. Wilkinson, Rajesh Odedra, Simon P. Heaton, Stephen R. Wedge, Nicholas J. Keen, Claire Crafter, John R. Foster, Madeleine C. Brady, Alison Bigley, Elaine Brown, Kate F. Byth, Nigel C. Barrass, Kirsten E. Mundt, Kevin M. Foote, Nicola M. Heron, Frederic H. Jung, Andrew A. Mortlock, F. Thomas Boyle, and Stephen Green. Azd1152, a selective inhibitor of aurora b kinase, inhibits human tumor xenograft growth by inducing apoptosis. *Clinical Cancer Research*, 13:3682–3688, 6 2007.

[30] Andrew A. Mortlock, Kevin M. Foote, Nicola M. Heron, Frédéric H. Jung, Georges Pasquet, Jean Jacques M. Lohmann, Nicolas Warin, Fabrice Renaud, Chris De Savi, Nicola J. Roberts, Trevor Johnson, Cyril B. Dousson, George B. Hill, David Perkins, Glenn Hatter, Robert W. Wilkinson, Stephen R. Wedge, Simon P. Heaton, Rajesh Odedra, Nicholas J. Keen, Claire Crafter, Elaine Brown, Katherine Thompson, Stephen Brightwell, Liz Khatri, Madeleine C. Brady, Sarah Kearney, David McKillop, Steve Rhead, Tony Parry, and Stephen Green. Discovery, synthesis, and in vivo activity of a new class of pyrazoloquinazolines as selective inhibitors of aurora b kinase. *Journal of Medicinal Chemistry*, 50:2213–2224, 5 2007.

[31] Hiroaki Ohno, Kazuo Kubo, Hideko Murooka, Yoshiko Kobayashi, Tsuyoshi Nishitoba, Masabumi Shibuya, Toshiyuki Yoneda, and Toshiyuki Isoe. A c-fms tyrosine kinase inhibitor, ki20227,suppresses osteoclast differentiation and osteolytic bone destruction in a bone metastasis model. *Molecular Cancer Therapeutics*, 5:2634–2643, 11 2006.

[32] Christopher W. Murray, Valerio Berdini, Ildiko M. Buck, Maria E. Carr, Anne Cleasby, Joseph E. Coyle, Jayne E. Curry, James E.H. Day, Phillip J. Day, Keisha Hearn, Aman Iqbal, Lydia Y.W. Lee, Vanessa Martins, Paul N. Mortenson, Joanne M. Munck, Lee W. Page, Sahil Patel, Susan Roomans, Kirsten Smith, Emiliano Tamanini, and Gordon Saxty. Fragment-based discovery of potent and selective ddr1/2 inhibitors. *ACS Medicinal Chemistry Letters*, 6:798–803, 7 2015.

[33] Ellen Weisberg, Paul W. Manley, Werner Breitenstein, Josef Brüggen, Sandra W. Cowan-Jacob, Arghya Ray, Brian Huntly, Doriano Fabbro, Gabriele Fendrich, Elizabeth Hall-Meyers, Andrew L. Kung, Jürgen

Mestan, George Q. Daley, Linda Callahan, Laurie Catley, Cara Cavazza, Azam Mohammed, Donna Neuberg, Renee D. Wright, D. Gary Gilliland, and James D. Griffin. Characterization of amn107, a selective inhibitor of native and mutant bcr-abl. *Cancer Cell*, 7:129–141, 2005.

[34] Malcolm P. Huestis, Matthew R. Durk, Charles Eigenbrot, Paul Gibbons, Thomas L. Hunsaker, Hank La, Dennis H. Leung, Wendy Liu, Shiva Malek, Mark Merchant, John G. Moffat, Christine S. Muli, Christine J. Orr, Brendan T. Parr, Frances Shanahan, Christopher J. Sneeringer, Weiru Wang, Ivana Yen, Jianping Yin, Joachim Rudolph, and Michael Siu. Targeting kras mutant cancers via combination treatment: Discovery of a pyridopyridazinone pan-raf kinase inhibitor. *ACS Medicinal Chemistry Letters*, 12:791–797, 5 2021.

[35] Ivana Yen, Frances Shanahan, Jeeyun Lee, Yong Sang Hong, Sang Joon Shin, Amanda R. Moore, Jawahar Sudhamsu, Matthew T. Chang, Inhwan Bae, Darlene Dela Cruz, Thomas Hunsaker, Christiaan Klijn, Nicholas P.D. Liau, Eva Lin, Scott E. Martin, Zora Modrusan, Robert Piskol, Ehud Segal, Avinashnarayan Venkatanarayan, Xin Ye, Jianping Yin, Liangxuan Zhang, Jin Soo Kim, Hyeong Seok Lim, Kyu Pyo Kim, Yu Jung Kim, Hye Sook Han, Soo Jung Lee, Seung Tae Kim, Minkyu Jung, Yoon hee Hong, Young Su Noh, Munjeong Choi, Oakpil Han, Malgorzata Nowicka, Shrividhya Srinivasan, Yibing Yan, Tae Won Kim, and Shiva Malek. Araf mutations confer resistance to the raf inhibitor belvarafenib in melanoma. *Nature*, 594:418–423, 6 2021.

[43] J rg Zimmermann, Elisabeth Buchdunger, Helmut Mett, Thomas Meyer, and Nicholas B Lydon. Potent and selective inhibitors of the abl-kinase: Phenylamino-pyrimidine (pap) derivatives. Technical report, 1997.

[52] Hans Richter, Alexander L. Satz, Marc Bedoucha, Bernd Buettelmann, Ann C. Petersen, Anja Harmeier, Ricardo Hermosilla, Remo Hochstrasser, Dominique Burger, Bernard Gsell, Rodolfo Gasser, Sylwia Huber, Melanie N. Hug, Buelent Kocer, Bernd Kuhn, Martin Ritter, Markus G. Rudolph, Franziska Weibel, Judith Molina-David, Jin Ju Kim, Javier Varona Santos, Martine Stihle, Guy J. Georges, R. Daniel Bonfil, Rafael Fridman, Sabine Uhles, Solange Moll, Christian Faul, Alessia Fornoni, and Marco Prunotto. Dna-encoded library-derived ddr1 inhibitor prevents fibrosis and renal function loss in a genetic mouse model of alport syndrome. *ACS Chemical Biology*, 14:37–49, 1 2019.

[53] Kumiko Nagashima, Stuart D. Shumway, Sriram Sathyanarayanan, Albert H. Chen, Brian Dolinski, Youyuan Xu, Heike Keilhack, Thi Nguyen, Maciej Wiznerowicz, Lixia Li, Bart A. Lutterbach, An Chi, Cloud Paweletz, Timothy Allison, Youwei Yan, Sanjeev K. Munshi, Anke Klippel, Manfred Kraus, Ekaterina V. Bobkova, Sujal Deshmukh, Zangwei Xu, Uwe Mueller, Alexander A. Szewczak, Bo Sheng Pan, Victoria Richon, Roy Pollock, Peter Blume-Jensen, Alan Northrup, and Jannik N. Andersen. Genetic and pharmacological inhibition of pdk1 in cancer cells: Characterization of a selective allosteric kinase inhibitor. *Journal of Biological Chemistry*, 286:6433–6448, 2 2011.

[54] Thomas Hanke, Sebastian Mathea, Julia Woortman, Eidarus Salah, Benedict Tilman Berger, Anthony Tumber, Risa Kashima, Akiko Hata, Bernhard Kuster, Susanne Müller, and Stefan Knapp. Development and characterization of type i, type ii, and type iii lim-kinase chemical probes. *Journal of Medicinal Chemistry*, 65:13264–13287, 10 2022.
